# A *topoietic* model of collective morphogenesis and compartmentalization

**DOI:** 10.64898/2026.11.03.763900

**Authors:** Santosh Manicka, Michael Levin

## Abstract

*Xenopus laevis* embryos survive teratogenic stress at significantly higher rates in larger groups, a phenomenon termed cross-embryo morphogenetic assistance (CEMA) (Tung, Sperry et al. 2024). We introduce *topoiesis*, a proposed mechanism by which a constitutively secreted stress signal (extracellular ATP or adenosine) diffuses and decays across the group, accumulating more in larger groups where interior volume outpaces absorbing surface. We find that this rising field carries more embryos to survival and demarcates the collective more sharply from its surroundings as collective screening strength grows with size. At intermediate sizes the outcome is spatially organized (a rescued interior enclosed by a failing periphery) and reconstitutes after shuffling. In larger groups the field restructures: its peak concentration keeps rising while its interior gradient flattens into a near-uniform bulk bounded by a sharp rim, sharpening demarcation while blurring distinctions among interior members. This pre-pattern stabilizes well before individual fate commitment, opening a window in which collective geometry can act on survival outcome. A biophysically calibrated minimal model capturing these principles recapitulates CEMA rescue rates across all six observed group sizes (*N* = 1–306; 0%–92%) and the size-dependent craniofacial defect rate among survivors. It predicts that sealing a group’s boundary against loss, rather than changing its size or biochemical regulation, moves mean rescue from 0% to 100% and reorganizes the rescue structure, tracking the predicted field geometry. These results reveal a hidden simplicity behind apparent complexity: diffusion and geometry alone may suffice to make a collective both spatially organized and controllable through macroscale intervention rather than microscale engineering of its members.

**Significance Statement:** Groups of animals can solve developmental problems that individuals cannot. We show that a simple physical mechanism, *topoiesis*, could explain how groups of *Xenopus laevis* embryos rescue one another from teratogenic stress at rates that climb steeply with group size. A constitutively secreted stress signal is lost to the surrounding medium, so larger groups retain more of it, a geometric principle shared with quorum sensing, nuclear criticality, and minimum viable patch size. The accumulating field that carries embryos to survival also sets the group apart from its surroundings, rendering larger collectives stronger. The environment that distinguishes a collective is therefore also the handle by which it can be steered, by shaping the group as a whole rather than its members.

## Introduction

Biological systems are often described as nested agents. Cells pursue small, local setpoints, such as keeping their voltage and pH within range, while collectives of cells pursue larger ones, such as building and repairing a specific body plan (Levin 2019, Levin 2022, McMillen and Levin 2024). Whether this nesting extends one level further, to groups of bodies, is not settled, and it raises a more basic question: how simple can the coupling between such bodies be, and still have consequences for the group as a whole?

The information that builds an embryo is usually traced vertically, from parent to offspring. Within the embryo, cells are thought to acquire positional information from a gradient, such as that emitted by a privileged organizer (a single, localized source), and to adopt their fates by threshold (Wolpert 1969, Crick 1970). In doing so, the embryo is treated as though it develops in isolation, its perimeter marking the natural boundary of morphogenesis. Yet embryos often develop alongside one another in a shared medium (whether a petridish, a clutch, or a uterus). Could a group of embryos, too, be a unit of morphogenesis, with information about position or fate available to its members? With no designated organizer among them, the simplest route is a shared signal: every embryo can emit it into the medium, and the information carried by the resulting concentration field depends on where the signal is lost and on whether its production depends on how much of it is already present.

In the extreme case where the signal stays within the shared volume and is spread evenly through it, as in a sealed or well-mixed culture, and every member emits, the concentration depends on the density of members, that is, on how many there are per unit volume. This is the basis of classical quorum sensing (Waters and Bassler 2005). A few cells confined in a small volume can therefore trigger the response that many cells trigger in a large one (Boedicker, Vincent et al. 2009), so the signal reports density rather than cell number. In this case the output is a single switch for the whole group, to which all members respond alike. Once signal can escape from the group, as in biofilms under flow, the concentration also depends on how quickly it is carried away and on how the cells are arranged in space (Redfield 2002, Hense, Kuttler et al. 2007), and the response then varies from place to place, being weaker where flow exposes cells and stronger where the signal is retained, for instance at the base of thick biofilms or inside crevices (Kim, Ingremeau et al. 2016).

A further example comes from ecology: a group with a leaky boundary, through which members are lost to the environment. If the group’s growth also depends on local density, there is a critical patch size below which it cannot persist. This result was derived independently for plankton blooms and for populations in fragmented habitat (Skellam 1951, Kierstead and Slobodkin 1953). Growth in proportion to local density is a positive feedback, since the population produces more of itself where it is already dense, and this feedback turns a graded profile into a sharp threshold on patch size. A developmental counterpart is the community effect, in which *Xenopus* muscle precursors commit to their fate only when a critical mass of like neighbours is present, through positive feedback among the cells (Gurdon 1988).

Whether any of these cases could allow embryos in a group to assist one another is a question about the limits of developmental robustness. Embryogenesis is buffered against genetic and environmental perturbation (Waddington 1942), but a teratogen above threshold overwhelms a single embryo. Ecology offers a precedent for groups faring better than individuals in the Allee effect, where individual fitness rises with population density (Allee 1931, Stephens, Sutherland et al. 1999). Mutual aid triggered by stress and carried through the medium has also been found in microbes: antibiotic-stressed *E. coli* release protein-carrying vesicles whose cell-free supernatant raises the survival of dormant neighbours in a concentration-dependent way (Wen, Bos et al. 2026), although through distinct donor and recipient states. If a comparable benefit operates among developing embryos that need not touch, it must travel through the shared medium, and embryos do influence that medium in ways other living systems can detect: nearby bacterial populations reorganize their spatial and ionic state, and the response follows the embryo when it is moved (Sergeeva, Pigozzi et al. 2026). The most plausible carrier is a fast, extracellularly diffusing cue. Extracellular ATP is a strong candidate: it is released on injury and stress, signals through P2 receptors across many tissues, and is degraded by ectonucleotidases within minutes (Burnstock 2007).

Such assistance has recently been quantified. *Xenopus laevis* embryos exposed to the teratogen thioridazine, at a constant dose and volume of medium per embryo, survive at rates that rise steeply and smoothly with group size, from 0% for singletons and groups of five to 13% at 25, 44% at 75, 85% at 100, and 98% at 300 (Tung, Sperry et al. 2024). No single group size marks a sharp onset. This effect, termed cross-embryo morphogenetic assistance (CEMA), is not specific to one compound. It also appears with forskolin and with expression of a dominant-negative Kir6.1 channel, where it shows up as graded craniofacial defect severity among survivors rather than as death. It was attributed to a short-range diffusible signal requiring calcium and P2 receptor signalling, and captured in an agent-based model of lateral interaction. Consistent with this, extracellular ATP measured directly in the medium was modestly higher in larger than in smaller groups under teratogen exposure, and no such difference was seen between untreated groups of the same sizes (Tung, Sperry et al. 2024). Because the volume of medium per embryo was held fixed, a culture in which the signal stays in the medium and is spread evenly through it would give the same signal at every group size. A dish keeps a non-volatile signal such as ATP in, but it need not mix it evenly: signal spreads from a group into the surrounding medium and decays along the way, so the amount a group retains can depend on its size. The rise is also smooth across the tested sizes, so it does not look like a feedback-driven threshold. This leaves the case in which every member emits at a constant rate, nothing feeds back on emission, and the medium outside the group absorbs the signal. Its physics is familiar from bacterial communities with leaky boundaries, but to our knowledge it has not been examined as an account of assistance among embryos, where it would have to turn group size into a graded, position-dependent chance of rescue. Can a group with no organizer, no privileged member and no perimeter of its own do this? Does that limit belong to its members, or does it emerge from the group’s interaction with its environment?

We suggest that this physics of a leaky boundary, familiar from the cases above, may also account for CEMA in a group of embryos. Suppose that every stressed embryo emits a signal at a constant rate, that the signal diffuses and decays, and that the medium outside the group absorbs it. The steady-state field is then not uniform. It is highest at the centre of the group, where the distance to the absorbing medium is greatest, and it falls toward the edge. Interior volume grows faster than absorbing surface, so larger groups retain a larger share of what they emit and the whole field rises with group size. We call this mechanism *topoiesis*. It requires only that the signal be emitted, diffuse, and decay, so the molecular identity of the carrier matters less than its physics. Two features distinguish it from the classical routes. The organizer is the group’s boundary, where the group meets the absorbing medium, rather than a cell, so the centre is simply the point farthest from that boundary and no molecular marker or privileged member is needed. Nor is the pattern a Turing instability, since the field is linear, with no interacting second species and no spontaneous symmetry breaking (Turing 1952, Gierer and Meinhardt 1972). The scale in this case is set by the ratio of group size to the signal’s decay length, *L*/*λ*, which determines how much of the group lies beyond the reach of the absorbing boundary and hence how the spatial character of the field changes as the group grows.

Here, we implement and analyze a minimal model of this mechanism, in which the coupling between embryos is nothing more than a shared, diffusing signal and an absorbing boundary at the edge of the group. It has four free parameters, two for the shared field and two for individual embryos, and the rest are anchored to independently measured biophysical quantities. The model reproduces the CEMA survival curve across the full range of group sizes. We do not tune its absolute timescales to observed ones, and the argument does not require us to. What matters is the separation between the timescales: the signal settles several-fold faster than individual fate commitments, so the field can be read as a pre-pattern rather than a readout. Topoiesis is the upstream half of the model. Downstream, each embryo’s own intra-embryonic morphogenesis is governed by a field-mediated bioelectric pre-patterning system that we introduced previously, which reproduces the qualitative developmental sequence of the craniofacial voltage pre-pattern characterized experimentally in *Xenopus* (Vandenberg, Morrie et al. 2011, Manicka and Levin 2025). The collective field shifts this system toward a normal or an aberrant developmental attractor (Methods), and an additional threshold on the deviation from the normal voltage pattern captures the group-size-dependent craniofacial defect rate among survivors. The model also makes predictions that the experiments have not yet tested. One is that the rescue signal itself is spatially structured within the group, highest at the centre and falling toward the edge. Another follows from the group fixing where its boundary lies while the medium fixes what happens there: acting on the shared medium around a group, rather than on its members, should be sufficient on its own. At a fixed group size, sealing the group’s boundary against loss moves mean rescue from 0% to 100%, monotonically in the fraction of the boundary sealed, without changing embryo number or chemistry.

## Results

### A minimal diffusion model reproduces group-size-dependent rescue

Why does simply being part of a larger group help a stressed embryo survive? Our model shows that a minimal mechanism, constitutive emission, diffusion and decay, and an absorbing boundary, could reproduce the full dependence of survival on group size with only four free parameters.

In our model, every stressed embryo constitutively emits a diffusible chemical signal, which we denote *F* (plausibly extracellular ATP or adenosine), into the shared medium; *F* diffuses and decays at rate *γ*F, and the group boundary is absorbing, fixing *F* to zero at the group edge (Fig. 1). The resulting field raises or fails to raise each embryo’s effective intracellular response past a fate-determining threshold, gated by a coupling gain *α* and each embryo’s intrinsic resilience *d*_0_; rescue is scored by whether the resulting voltage pattern falls within a criterion distance (mean squared deviation, MSD) of the normal target pattern (full derivation and criterion justification in Methods). The model, therefore, has four free parameters: the field decay rate *γ*F and coupling gain *α*, fit together with the spread of intrinsic resilience *σ* and the MSD rescue threshold so that the model reproduces the observed CEMA rescue curve. The remaining parameters, including diffusivity, embryo spacing, and peak emission rate, are either anchored to independently measured biophysical quantities or assumed a priori (calibration table in Methods).

**Figure 1.**
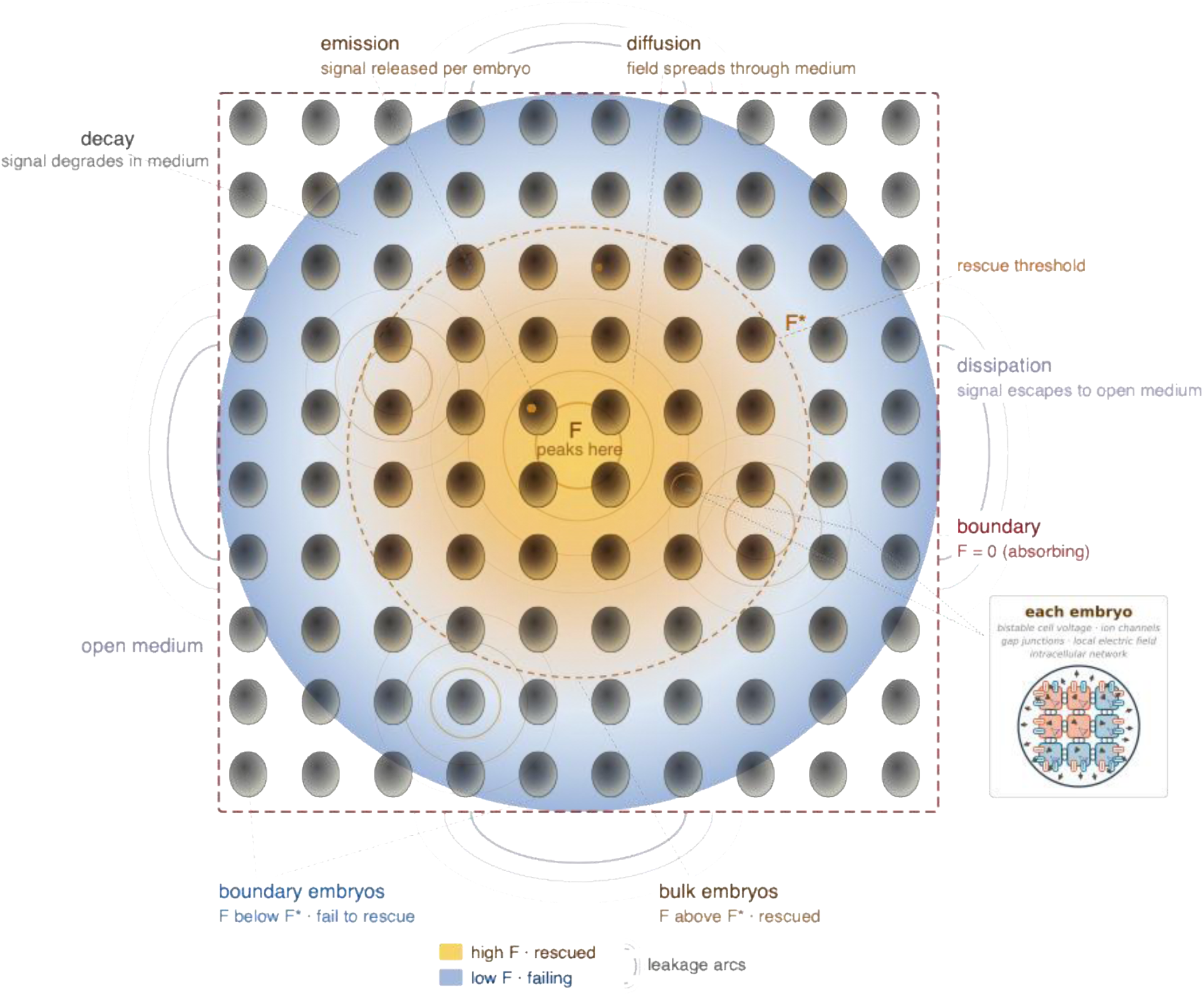
The topoiesis mechanism. A collective of stressed Xenopus laevis embryos constitutively emits a signal *F* (eATP/adenosine) that diffuses through the shared extracellular medium and decays at rate *γF*. Absorbing boundary conditions at the group edge (open medium; *F* = 0) impose a geometric asymmetry: boundary embryos lose field rapidly via leakage arcs, while bulk embryos are shielded by their neighbors. At steady state *FSS* peaks at the group center and decays monotonically to zero at the periphery. Embryos above the rescue threshold (bulk; dashed contour) are rescued; those below (boundary) are not. Each embryo is itself a bioelectric tissue: a lattice of bistable cells coupled by voltage-gated gap junctions, enmeshed in an intra-embryonic electric field that both arises from their membrane potentials and regulates them (Manicka and Levin 2025), with an intracellular biochemical network in each cell setting how strongly that cell responds to the field (inset). The collective signal *F* acts on this machinery by shifting each embryo toward the normal or the aberrant bioelectric pre-pattern (Methods).

This minimal model reproduces the CEMA survival curve across the observed range of group sizes (Fig. 2a). A single additional threshold, applied only within the population the model already scores as rescued, captures the group-size-dependent craniofacial defect rate reported for a second, unrelated perturbation (Kir6.1 dominant-negative) among survivors, with no re-fitting of *γ*F or *α* (Fig. 2b). The settled bioelectric pattern underlying each rescue outcome and defect classification, for a representative low-rescue and a representative high-rescue group, is shown in Fig. 3.

**Figure 2.**
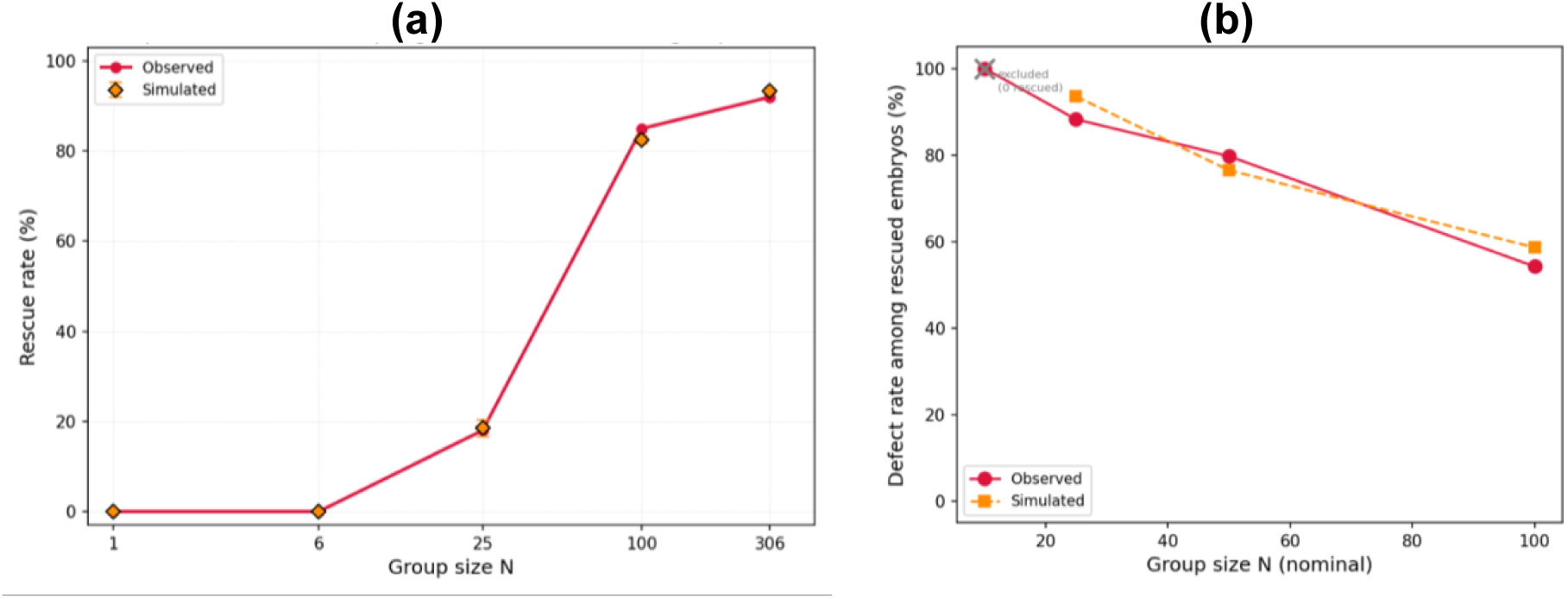
The model reproduces CEMA rescue rates and, with one additional threshold, an independent defect-rate curve. (a) Observed rescue rates (red circles; (Tung, Sperry et al. 2024)) and simulated rates (orange diamonds; canonical parameters *α* = 400, *γF* = 1.845 × 10^−4^ s^−1^, *D*0 = 0.01, *σ* = 0.040) across six group sizes (*N* = 1–306). The model reproduces the sigmoidal rescue transition with no rescue at *N* ≤ 6 and > 80% rescue at *N* ≥ 100. **(b)** Simulated (orange) versus observed (red) craniofacial defect rate among survivors of a Kir6.1 dominant-negative perturbation, across *N* ∈ {10,25,50,100} (*N* = 10 excluded: below the critical rescue size, so no survivors are scored). A single fitted threshold, applied post hoc to the already-rescued population from panel (a), matches the observed curve within ±5.3 percentage points.

**Figure 3.**
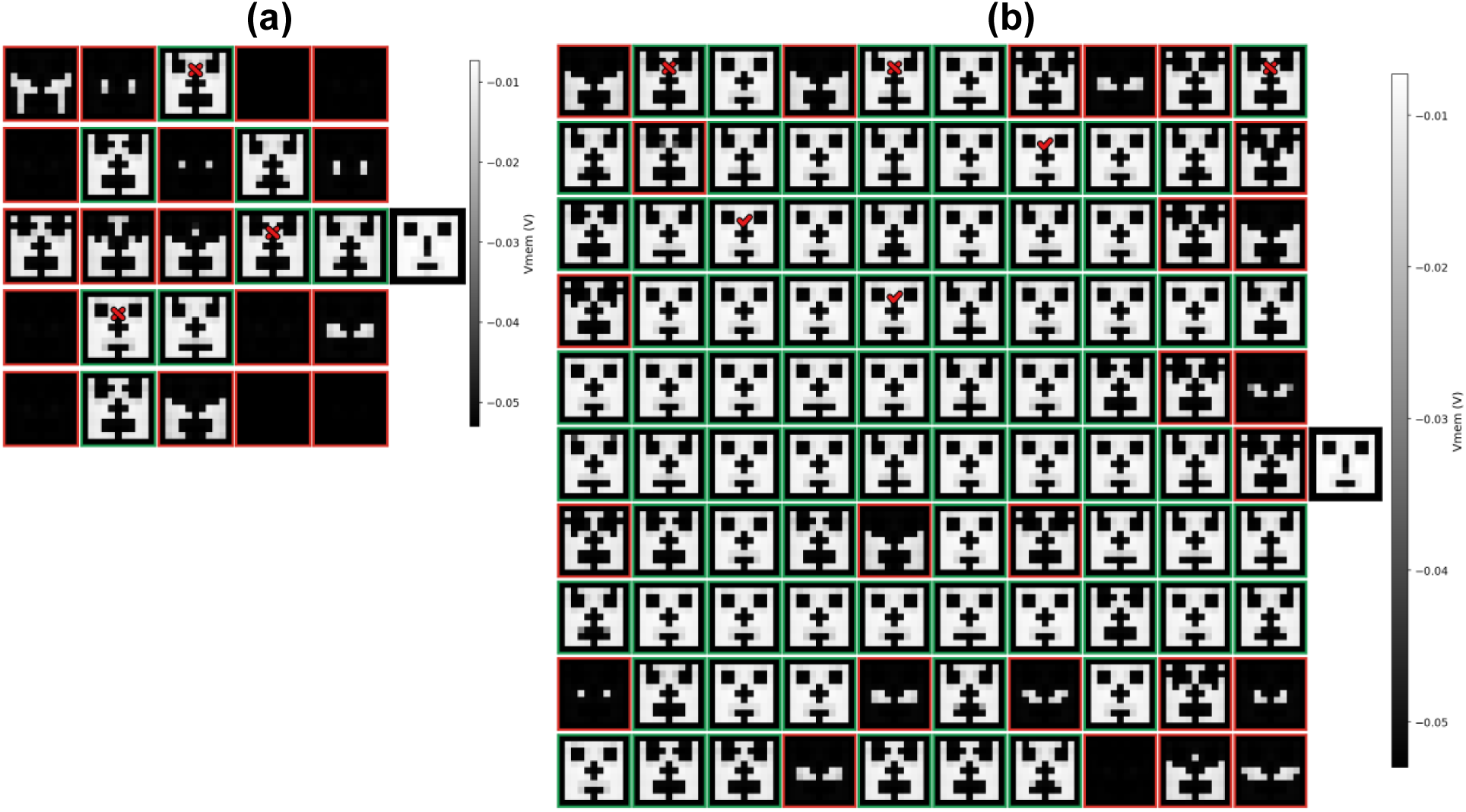
The settled bioelectric pattern of every embryo in a representative trial, corresponding to the low– and high-rescue conditions in Fig. 2a. Each panel is one embryo’s final Vmem pattern; green border indicates rescue, red indicates failure, and black marks the reference pattern. Red checkmarks and crosses mark several non-adjacent representative “normal” and “defective” examples (the categories underlying the craniofacial defect-rate threshold in Fig. 2b), spread across the grid rather than confined to one region. Trials shown were chosen to display both rescue outcomes clearly and are not the population average; Fig. 2a reports the trial-averaged curve. **(a)** *N* = 25, 8/25 rescued. No rescued embryo in this trial falls below the defect threshold, so no “normal” examples are marked. **(b)** *N* = 100, 75/100 rescued, with both categories represented.

### A spatially structured stress field forms before fate commitment and pre-patterns the rescue map

Within a stressed group of embryos, does an embryo’s position predict whether it survives, and is that positional signature already present before its fate is decided? Our model produces a field that reaches its steady-state spatial pattern several-fold before each embryo’s intracellular machinery commits to a fate, and rescue outcomes averaged over many trials closely track this pre-existing pattern.

In our simulations, *F* reaches steady state within roughly 48 minutes of stress onset, about 3.5 times faster than the intracellular machinery commits to a fate, which occurs only around 2.8 hours later (Fig. 4). Because the field’s spatial pattern is already fixed well before any embryo’s fate is decided, it functions as a genuine pre-pattern: collective geometry can act as a cause of the ensuing rescue outcome, not merely a correlate of it.

**Figure 4.**
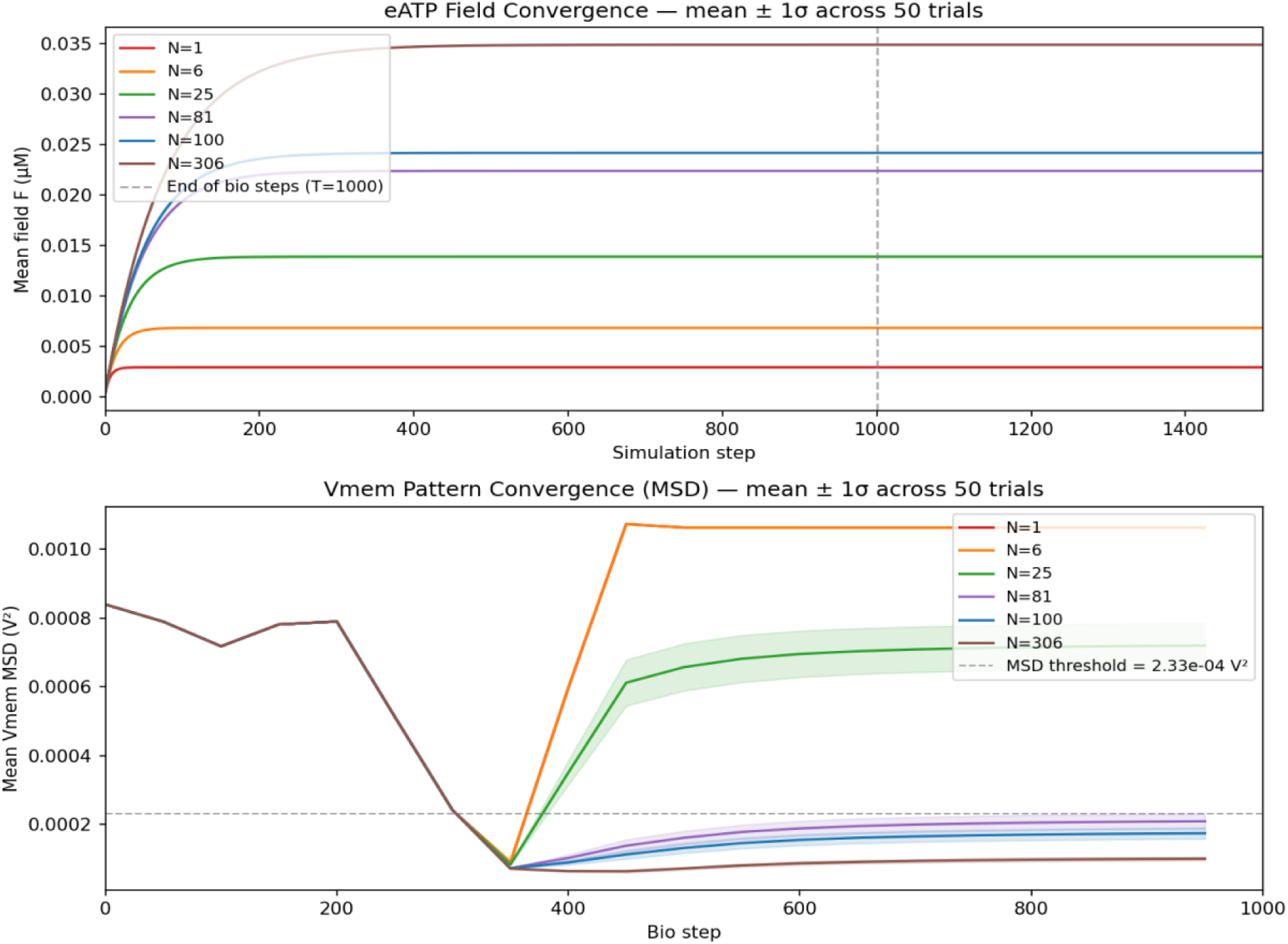
The field reaches steady state well before fate commits. **Top:** mean field *F* over simulation time for each group size (mean ± 1 s.d., 50 trials). *F* saturates within roughly the first 100–300 steps at every *N*, corresponding to ∼48 min of biological time under the timestep calibration in Methods. **Bottom:** mean V_mem_ pattern deviation (MSD) over the same runs. Curves for all group sizes remain superimposed until ∼ step 350, corresponding to ∼2.8 h: the shared early dip is whole-pattern transient recovery present in every embryo regardless of eventual outcome, and the divergence that follows, rescued patterns retaining this recovered face while failing ones disintegrate, is driven by a saddle-node bifurcation in the interior cells’ polarising-channel conductance (SI Appendix 4). The field’s spatial pattern is therefore fixed several-fold earlier than the outcome it predicts.

At steady state, *F* is highest at the group centre and falls toward the boundary at every group size we simulated (Fig. 5, middle row), with the centre-to-corner gradient growing from 61% at *N* = 25 to 79% at *N* = 100 as more of the group is carried above the approximate field concentration needed for rescue (SI Appendix 6). A closed-form solution for this centre-field growth is derived in Methods; it reproduces the correct qualitative scaling but not the absolute magnitude (SI Appendix 1). By *N* = 210–306 the interior has flattened into a broad, near-uniform plateau, a change in the field’s character we quantify in the next section. An embryo’s intrinsic resilience *d*_0_, its capacity to reach the normal pattern without help from the collective, carries no spatial structure (Fig. 5, top row): each embryo draws its value independently from a probability distribution (Methods), so a trial-averaged map of *d*_0_ across the group is uniform noise by construction. Field and resilience contribute additively to determine each embryo’s fate (Methods), so this asymmetry, a highly structured field against an unstructured resilience, raises the question of which of the two determines an embryo’s fate.

**Figure 5.**
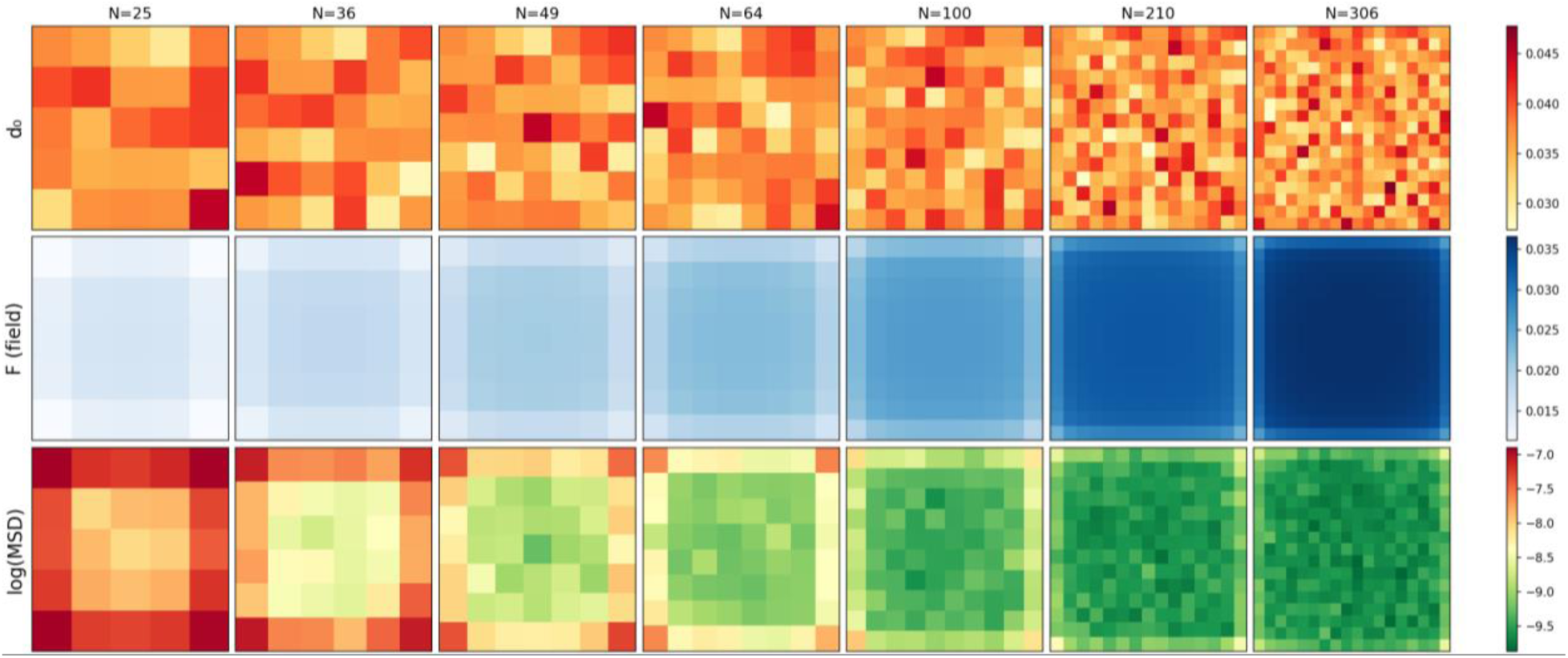
Trial-averaged maps show a highly structured field, an unstructured intrinsic resilience, and a rescue outcome that tracks the field. Each column is one group size (*N* = 25 to 306); each panel is averaged over 50 trials. **Top:** intrinsic resilience *d*_0_, drawn independently per embryo per trial, shows no spatial structure at any *N*. **Middle:** steady-state field *F* is highest at the group centre and falls toward the boundary; the gradient is steep at small *N* and progressively flattens into a broad interior plateau by *N* = 210–306. **Bottom:** *log*(*MSD*) (rescue outcome; lower is better-rescued) mirrors the field’s spatial structure at every group size, most sharply where the field itself is steepest.

Its immediate consequence for the rescue map is that any single trial inherits the variation in *d*_0_ rather than reflecting *F* alone. At *N* = 25, near threshold, a single trial’s outcome appears spatially disorganised, with little evident relation between position and rescue; at *N* = 100, well above threshold, a single trial already shows a clear centre-rescued pattern, because enough of the group exceeds the level needed for rescue by a sufficient margin that *d*_0_ rarely reverses an embryo’s outcome (single-trial maps, SI Appendix 2). Averaging over trials cancels the variation in *d*_0_: the resulting pattern (Fig. 5, bottom row) tracks *F* closely at every group size we tested (trial-averaged *r*^2^ = 0.72–0.94), recovering a centre-to-edge contrast that any single trial obscures, at *N* = 25 a span of roughly 0.4 in rescue probability across the group, and at *N* = 100 near-universal interior rescue with only corner-confined failures.

A direct causal test confirms that the field is responsible for this pattern rather than merely correlated with it. Holding *d*_0_ fixed and forcing *F* to a chosen value in an isolated embryo, a two-way analysis of variance attributes 79–82% of the variance in outcome to *F* and 6.4% to *d*_0_; at either extreme of the range tested, *F* alone determines the outcome irrespective of *d*_0_, whereas no value of *d*_0_ does so irrespective of *F* (SI Appendix).

### The field compartmentalizes the collective more sharply with size, and acts on position rather than identity within it

Do embryonic collectives attain a group-level identity of their own (a boundary that sets them apart from their surroundings), and does that identity sharpen as the collective grows? Our model predicts that they should: as a group grows, the contrast between the field concentration at its centre and at its boundary grows, though at the cost of precision about where, within it, any single embryo sits, and the spatial pattern it imposes on its members is a real, re-impressible template rather than a fixed record of who they are.

The field’s two roles, demarcating the group from its surroundings and resolving position within it, scale in opposite directions with group size (Fig. 6). The per-cell steepness of its gradient, which is what allows neighbouring positions to be distinguished, falls roughly fourfold from *N* = 25 to *N* = 306, and the trial-averaged tracking established above becomes correspondingly less precise at large *N*: *r*^2^ falls from 0.94 to 0.72 over the same range. The contrast between the group’s interior and its boundary moves in the opposite direction, growing about twofold. Both trends follow from the same underlying fact: the environment keeps a small group below the concentration it would otherwise reach, while a large group increasingly approaches that concentration unconstrained, not because it behaves differently, but because growing the group increases the fraction of its members that already sit far enough from the boundary to be effectively beyond its reach (Eqs. 4, 5). Near the boundary, an embryo’s field is held below the concentration emission and decay alone would set; beyond a few decay lengths *λ* from the boundary, it reaches that concentration regardless of where the boundary lies. The centre-to-boundary contrast therefore grows because more of the interior occupies this unconstrained region, while the boundary’s own pull stays exactly as strong and confined to a rim of fixed width ∼ *λ* rather than widening with the group. Spread across a progressively larger domain, that same transition yields a shallower per-cell slope. Larger groups therefore build a sharper boundary against their surroundings even as they resolve less detail within it, and the rescued-centre, failing-periphery body plan is crispest at intermediate group sizes and washes out, though rescue itself does not, once the group is large.

**Figure 6.**
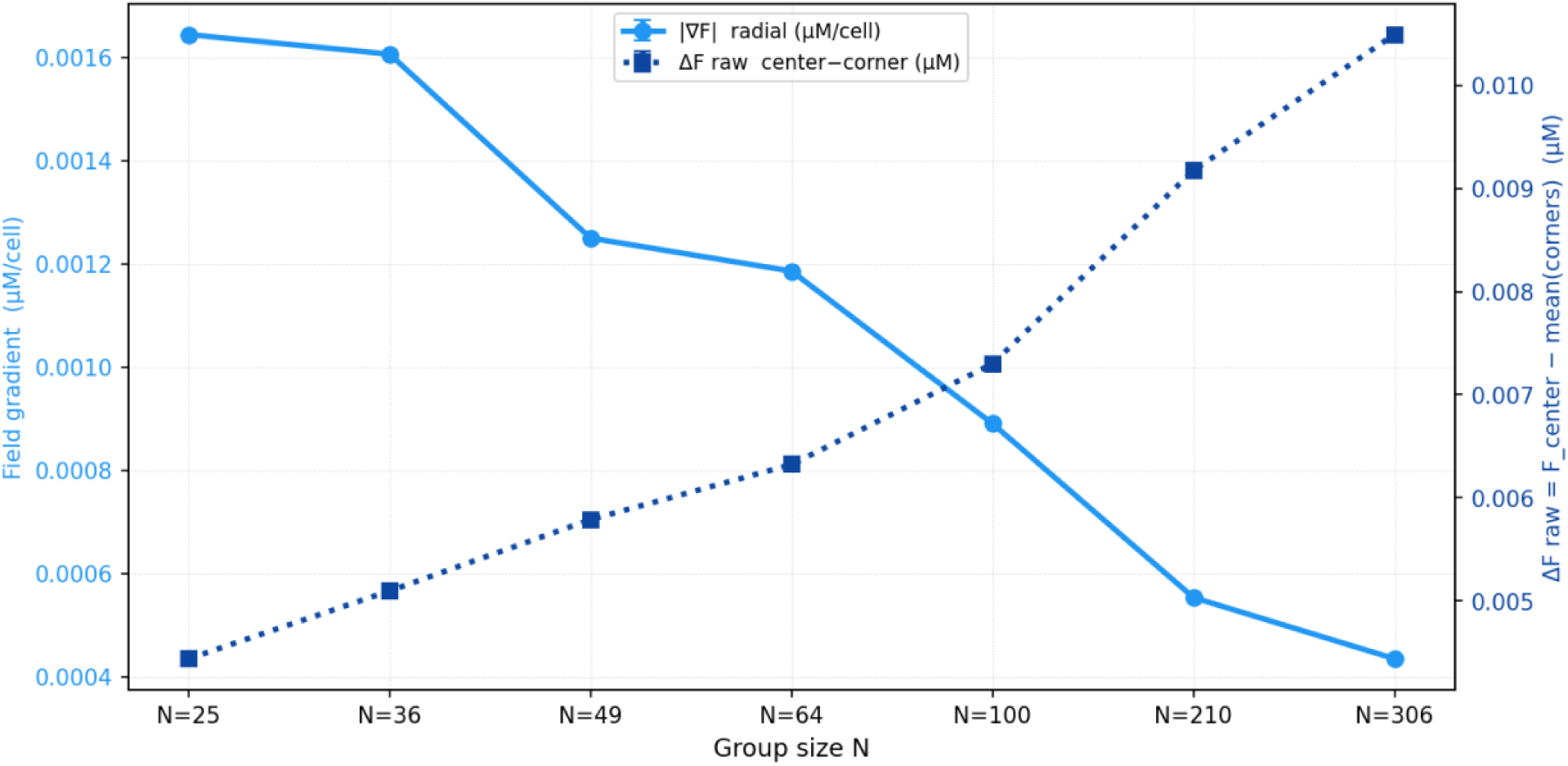
Compartmentalization of the group strengthens while resolution of position within it weakens, as the group grows. Both quantities are averaged over 50 trials per group size. The per-cell gradient |*∇F*| (left axis, solid), which sets how finely neighbouring positions can be distinguished, falls roughly fourfold from *N* = 25 to *N* = 306. The raw centre-to-corner contrast *ΔF* (right axis, dotted; unsmoothed), which sets how sharply the group as a whole stands out against its surroundings, grows about twofold over the same range. Larger groups therefore demarcate themselves more strongly while resolving position within themselves less precisely.

A direct perturbation establishes that this spatial pattern is imposed by the field on whichever embryos occupy each position, rather than being a record of outcomes already assigned. We relocated embryos to new positions within the group at a chosen time after stress onset and let the simulation continue, which dissociates position from identity: averaging the resulting rescue map over many trials and comparing it against the trial-averaged unshuffled map indicates whether the ensemble-level spatial pattern follows the field at each position or instead follows the relocated embryos. Because a trial-averaged map of finite size is itself only reproducible up to a noise floor, and that floor rises with *N* as the field flattens, we express each comparison relative to a per-*N* ceiling, the agreement two independent trial-averaged baselines achieve with one another (Methods). A ratio of 1.0 denotes reconstitution as complete as the data permit.

Early shuffles reconstitute the map fully at every group size. For *t* ≤ 20 bio-steps after stress onset the ratio sits at the ceiling throughout the range tested, from *N* = 25 to *N* = 306 (Fig. 7): the field re-imposes the same spatial pattern on whichever embryos now occupy each position, and it does so no less completely in large groups than in small ones. What declines with *N* is the ceiling itself, from *r*^2^ = 0.89 at *N* = 25 to 0.56 at *N* = 306. This follows from the flattening gradient of Fig. 6: the position-to-position differences that distinguish one map from another shrink as |∇*F*| falls, while the per-embryo variation in *d*_0_ that obscures them does not change with group size, so averaging a fixed number of trials recovers a progressively smaller fraction of the underlying map. In other words, the template becomes less reproducible without becoming less faithfully re-imposed. Late shuffles behave differently. From *t* = 50 onward the ratio falls below the ceiling, and the shortfall deepens with group size: at *t* = 100 it remains at the ceiling for *N* = 25 but falls to 0.04 at *N* = 306. This is genuine erasure by commitment rather than an effect of the noise floor, since it survives normalisation. Together the two regimes bracket a positional commitment window, within which an embryo’s fate can still be redirected by where it ends up and beyond which it cannot. The collective’s spatial pattern is therefore transiently rewritable: after an early displacement the same positions are rescued again, while which particular embryos are rescued continues to track their own resilience rather than their new location (SI Appendix 5).

**Figure 7.**
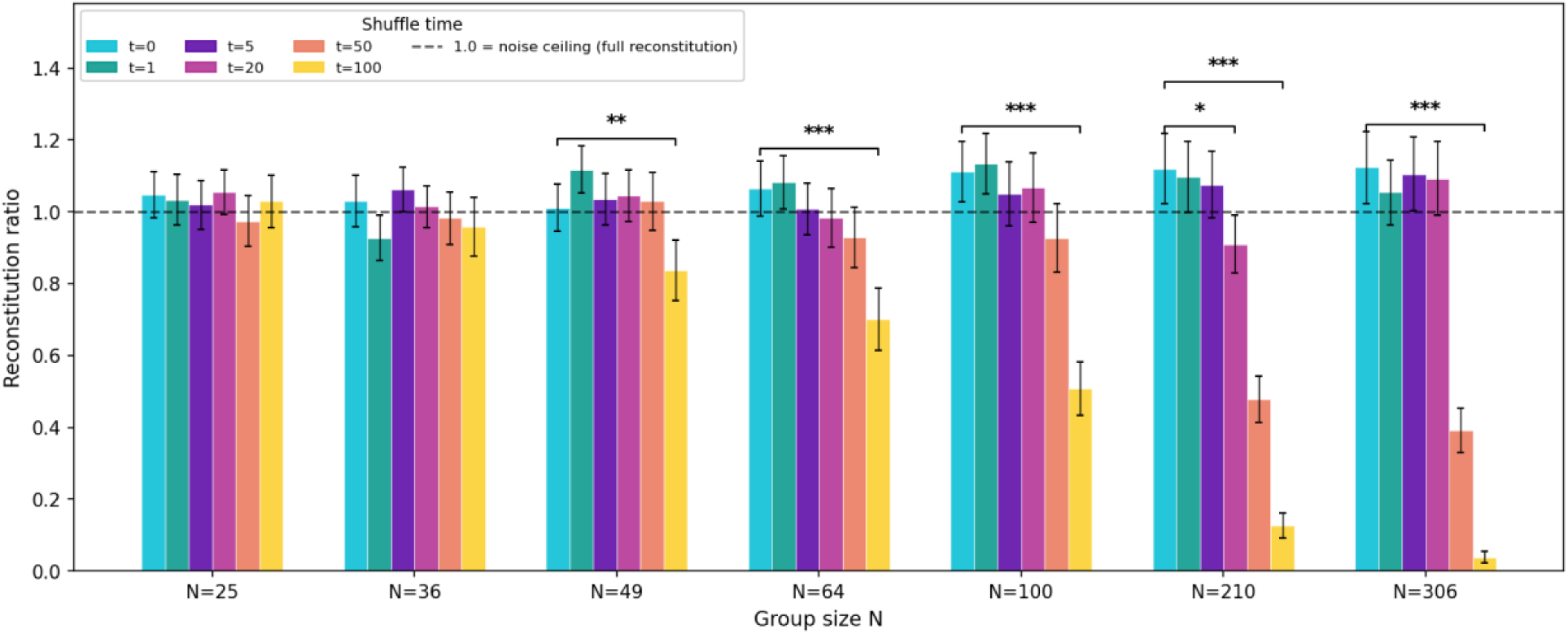
Early shuffles reconstitute the spatial pattern fully at every group size; late shuffles erase it, increasingly so as the group grows. Agreement (Pearson *r*^2^) between the trial-averaged rescue map of a shuffle-at-time-*t* run and the trial-averaged unshuffled baseline, divided by a per-*N* noise ceiling: the agreement two independent trial-averaged baselines achieve with one another, obtained by splitting the 50 baseline trials into halves and applying the Spearman–Brown correction (Methods). A ratio of 1.0, marked by the dashed line, denotes reconstitution as complete as the data permit. Bars are ratios with bootstrap standard errors, across group size and shuffle time (*t* ∈ {0,1,5,20,50,100} bio-steps). For *t* ≤ 20 the ratio sits at the ceiling at every *N*; from *t* = 50 onward it falls below, and the shortfall deepens with group size, reaching 0.04 at *N* = 306 by *t* = 100. Asterisks mark significant differences from t=0 within the same N (∗q<0.05, ∗∗q<0.01, ∗∗∗q<0.001; paired bootstrap test, Benjamini–Hochberg corrected across all fourteen N × {t=20, 100} tests; Methods): the t=100 shortfall is significant from N=49 onward, while t=20 differs significantly from t=0 only at N=210. Ratios slightly above one arise because the shuffled runs reuse the baseline field realisation, so the comparison shares a source of variance that the disjoint-halves ceiling does not; this makes the ceiling a conservative reference rather than an inflated one. The figure reports the positional component of the shuffle result; the identity component is a separate metric, reported numerically in the main text.

### Sealing the group’s boundary controls rescue magnitude and steers its spatial pattern

Can the medium surrounding a group be used to redirect and amplify rescue without altering embryo number or chemistry? Our model predicts that it can: at a fixed group size, sealing the boundary against loss moves mean rescue from failure to saturation, and which embryos within the group are rescued tracks which walls are sealed.

We tested five boundary conditions at *N* = 25 (10 trials each), all absorbing except for zero or more walls made reflecting: the default fully absorbing boundary, one reflecting wall (north), two reflecting walls (adjacent, north+west, or opposite, north+south), and all four reflecting. Mean rescue rises monotonically with the number of sealed walls: 0% (absorbing) → 21.6% (one wall) →∼ 76% (two walls, 76.0% adjacent and 75.6% opposite, the two configurations statistically indistinguishable) → 100% (all four); Fig. 8 shows the two extremes and the one-wall case, the three conditions that most clearly illustrate the trend. The same 25 embryos, the same coupling gain and decay rate, the same emission: the full range of the observed rescue curve, from complete failure to complete rescue, is reachable from boundary conditions alone at a group size fixed throughout.

**Figure 8.**
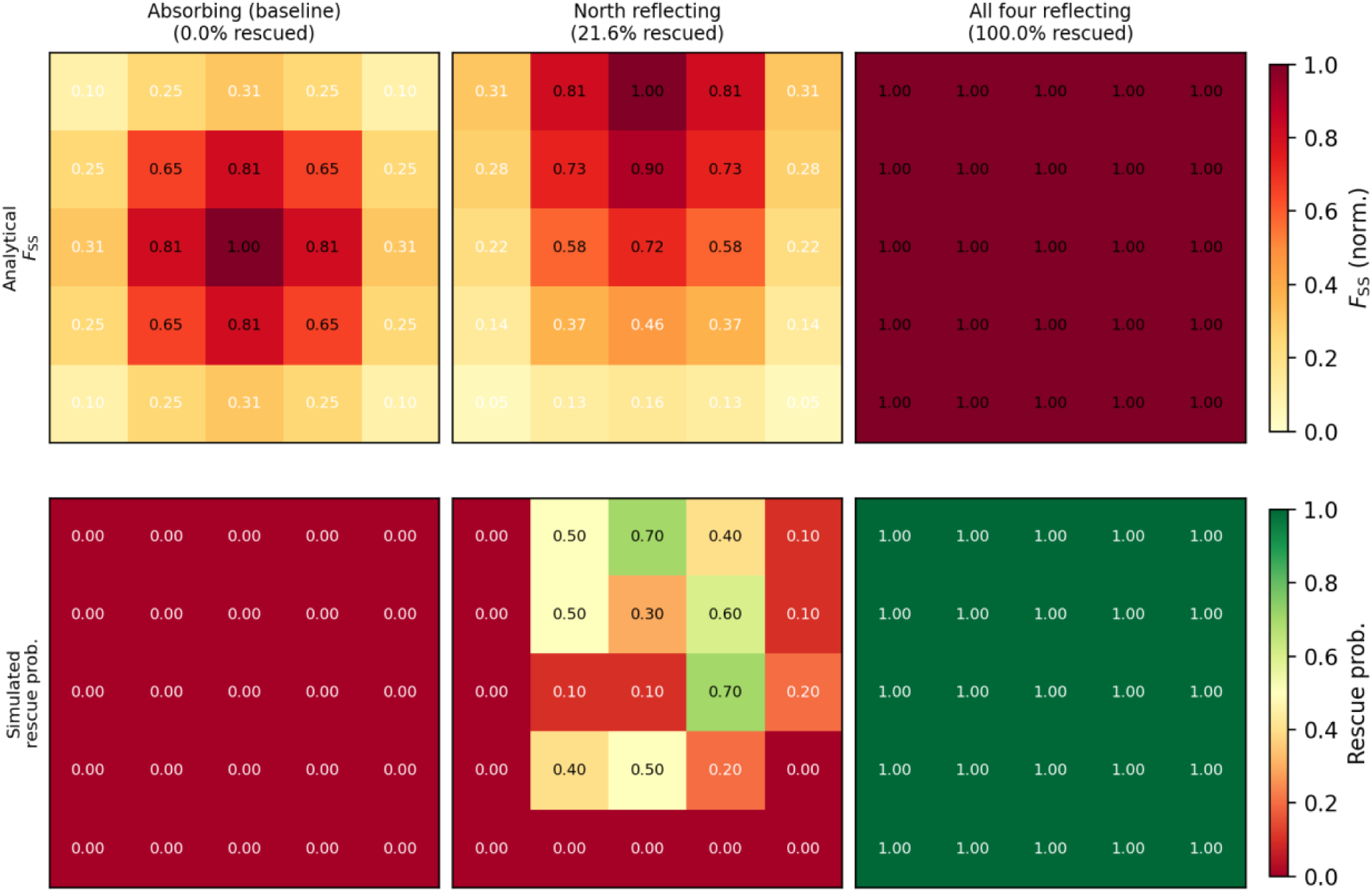
Sealing the group’s boundary against loss raises rescue and relocates it, tracking the analytically predicted field eigenmode. *N* = 25, 10 trials per condition. **Top:** analytical steady-state field eigenmode *FSS* (normalised), absorbing boundaries (centre-peaked), one reflecting wall (north; peak shifted to the top rows), and all four walls reflecting (spatially uniform). **Bottom:** trial-averaged simulated rescue probability at each grid position, tracking the eigenmode above it. Column titles give the mean rescue rate across all 25 positions and 10 trials. Two intermediate configurations (two reflecting walls, adjacent or opposite) are omitted from the figure; both give ≈ 76% mean rescue (main text), consistent with shifting the eigenmode peak off-centre by a similar amount along different axes.

Sealing a wall does not merely raise the rescue rate; it relocates where rescue occurs, tracking the analytically predicted field eigenmode for each configuration. Absorbing boundaries give the familiar centre-peaked mode and, at this group size, too little rescue to score; sealing the north wall shifts the peak to the top rows, and the trial-averaged rescue map follows it there (Fig. 8, comparing the analytical and simulated rows). The spatial pattern of rescue is therefore not a fixed property of the group. It is read off the field’s geometry, and that geometry is set jointly by which walls leak.

Sealing all four walls removes boundary loss entirely, so the steady state is uniform at the ceiling set by emission and decay alone, *F* = *P*/*γ*F (Methods): every embryo receives the same field regardless of position, and all are rescued. This is the same limit a sufficiently large absorbing-boundary group approaches from within: at the group centre, *F*SS(0) = (*P*/*γ*F)[1 − sech(*L*/2*λ*)] (Eq. 4), and the same sech(*L*/2*λ*) term that vanishes as *L* grows is set to zero directly, at any group size, by removing the boundary that produces it. Sealing the boundary of a small group therefore reaches, at fixed *N*, the same field homogeneity that a much larger absorbing-boundary group reaches only by growing past its edge. Boundary control and group-size increase are two routes to the same outcome, which makes the group’s boundary condition not merely a modelling assumption but an experimentally addressable lever on collective behavior.

## Discussion

In this work we asked whether a single diffusible signal, emitted constitutively and lost to the surrounding medium, could account for the group-size dependence of teratogen resistance in systems such as embryonic *Xenopus*, and our model predicts that it can. Because the field this signal forms is established well before the embryo’s morphogenetic pattern commits, it can act as a cause of the rescue outcome rather than a correlate of it, and the rescue map that follows tracks it closely once individual variation is averaged away. Holding resilience fixed and forcing the field directly confirmed the direction of that dependence: the field, rather than an embryo’s own resilience, determines the outcome. Because the same field serves two roles at once, growth improves one at the expense of the other, widening the contrast between the field’s concentration at its center and at its boundary while resolving position within it less precisely, so that the spatial body plan is sharpest at intermediate sizes. What that pattern amounts to is a template the field re-imposes on whichever embryos occupy each position, rather than a record of fates already assigned.

Our model and analysis suggest two experiments that could test the mechanism directly, and both act on the medium rather than on the embryos. First, lowering the signal’s degradation rate by ectonucleotidase inhibition, or raising its ambient level by adding extracellular ATP, should rescue groups that lie below the critical size, since both raise the interior field without altering the number of embryos or their spacing. Second, rescue should depend on the shape of the domain and not only on its area: a compact group screens its interior more effectively than an elongated one of equal number, so rescue rate should fall as a well is made more eccentric, and a single-file arrangement should not rescue at any group size. Both predictions follow from the same premise, that what a collective can do is set by the geometry of the medium it occupies, and both are the experimental form of the claim that such collectives can be steered as wholes by shaping their surroundings.

What connects teratogen exposure to the connectivity modulation our model applies to each embryo’s network? A plausible chain could run as follows. Thioridazine itself activates calcium and reactive-oxygen signalling directly, though this has so far only been shown in heart cells, not embryos (Liu, Xu et al. 2020). Calcium-linked stress is a well-established trigger for ATP release through connexin and pannexin hemichannels (Bao, Locovei et al. 2004). Once released, ATP and its breakdown product adenosine diffuse and decay on a timescale typical of purinergic signalling generally (Burnstock 2007). *Xenopus* embryos express the relevant purinergic receptors and ectonucleotidases at early developmental stages (Blanchard, Boué-Grabot et al. 2019, Massé, Bhamra et al. 2021), and purinergic signalling plays a broad role in vertebrate embryonic development more generally (Burnstock and Ulrich 2011). Downstream, receptors of this kind often act through kinase cascades that behave as switches. CaMKII’s autophosphorylating bistability is the classic example (Lisman 1985): a continuous biochemical state that behaves as a switch because of its own positive feedback. Effective damping (Eq. 6) does the same in our model: it continuously scales the network’s weights, and the network’s own bistability turns that scaling into a threshold crossing, as CaMKII’s does. Tung et al. implicated calcium and P2 receptor signalling in CEMA, but found that chelating intracellular calcium did not block the effect (Tung, Sperry et al. 2024). Our model does not need to explain this away, since emission here is gated by a stress variable with no calcium dependence of its own (Eq. 1). Two points in this chain still remain open, namely, which receptor and kinase are responsible, and what a given teratogen does at the cellular level to trigger release in the first place. We offer this as a plausibility case for the kind of mechanism the model requires, not a claim to have identified the actual molecules.

That plausibility case has an obvious limitation: it targets one particular chain among several that could in principle connect teratogen exposure to rescue, and leaves the molecular identity of every step underdetermined. Minimal models with exactly this kind of limitation are the norm in this genre, however, not the exception. Hodgkin and Huxley’s account of the action potential was predictive for years before the identity of the channels it described was known (Hodgkin and Huxley 1952), and neither the activator-inhibitor route (Turing 1952) nor its explicit reaction-diffusion form (Gierer and Meinhardt 1972) specifies what those species are biochemically. Quorum sensing offers a direct empirical precedent for why this move is safe: confining as few as one to three bacteria in a small enough volume triggers the same gene expression that thousands trigger in an open one (Boedicker, Vincent et al. 2009), showing that what launches the response is local concentration, jointly set by population and geometry, not a count of cells. The elaborate molecular machinery of quorum sensing operates downstream of that trigger, not as part of it. Our model targets the analogous upstream layer, the geometry that sets a diffusible signal’s concentration, and leaves what happens once threshold is crossed, addressed above for this system specifically, as a separate question.

These results place the collective between an aggregate and an organism. The rescue signal field is a morphogenetic field in the sense originally developed by Driesch, Gurwitsch and Weiss and formalised since (French, Bryant et al. 1976, Goodwin and Trainor 1980, De Robertis 2009, Levin 2011): spatially ordered, regulative in that it reconstitutes after displacement, a property of the ensemble rather than of any member, and organised about a centre fixed by geometry rather than by a molecular marker. The collective does not, however, produce its own boundary. It generates the field’s amplitude through a closed loop of emission and response, but the zero against which that amplitude is read is imposed by the absorbing environment. The collective therefore fixes where its boundary lies while the medium fixes what happens there, which makes it a “pre-closure” individual whose coherence is scaffolded by its surroundings rather than self-produced (Varela, Maturana et al. 1974, Black, Bourrat et al. 2020). That a system’s individuality can be graded by how much of its organisation it supplies itself is not a new idea (Levin 2019, Krakauer, Bertschinger et al. 2020); what topoiesis contributes is a case in which the grading is a boundary condition on a diffusion equation with measurable consequences, and a demarcating mechanism that is field-borne rather than structural, which adds an entry to the catalogue of ways individuality can be realised (Clarke 2013).

The governing quantity, the ratio of group size to the signal’s decay length, recurs wherever something is produced throughout a volume and lost at its boundary. It sets the minimum patch size for a plankton bloom or a population in fragmented habitat (Skellam 1951, Kierstead and Slobodkin 1953), the critical mass for neutron multiplication, and the size at which a huddle of penguins retains more heat than it sheds. Quorum sensing is the closest biological relative, and the reading on which a shared diffusible signal reports geometry and mass transfer rather than headcount (Redfield 2002, Hense, Kuttler et al. 2007) is what topoiesis makes explicit, with the addition that an absorbing boundary converts that report into a spatial coordinate. In ecology the corresponding observation is the Allee effect, in which individual fitness rises with population density (Allee 1931, Stephens, Sutherland et al. 1999).

Taken together, the result is that a collective acquires both a boundary and an internal coordinate from nothing more than a shared signal and a lossy environment. Supra-organismal order of this kind requires no dedicated machinery and may therefore arise wherever organisms develop in a shared medium, which would make the cohort, rather than the individual, the appropriate unit for developmental and toxicological experiments, and perhaps for synthetic bioengineering approaches that could capitalize on these effects for desired properties of robustness and functionality.

## Materials and Methods

### Model description

The model has two levels: machinery within each embryo that patterns its tissue, and a signal shared between embryos that couples them into a collective.

Teratogen exposure raises each embryo’s stress. This process is autonomous, independent of the embryo’s bioelectric state: stress is self-reinforcing once triggered and opposed by a fixed decay, so an embryo settles into either a low, unstressed state or a high, sustained one, rather than tracking the teratogen level continuously. Every stressed embryo constitutively emits a diffusible signal, in proportion to its own current stress level, into the shared medium. Unstressed embryos emit nothing, consistent with the finding that untreated embryos reared with treated ones do not aid them (Tung, Sperry et al. 2024). The signal diffuses, decays at a first-order rate, and is absorbed at the boundary of the group, where the surrounding medium holds it at zero. Because interior volume grows faster than absorbing surface, larger groups retain more of what they emit, and the resulting steady-state field is highest at the group centre and falls toward the edge.

This shared field acts back on every embryo throughout the run by shifting the effective damping of its intracellular biochemical network: the field and the embryo’s own intrinsic resilience, an independently assigned baseline capacity to reach the normal developmental program without the field’s help, combine to set this shift, scaling how strongly the network’s own dynamics respond and, through it, each cell’s sensitivity to the local field described below.

The biochemical network so scaled operates within the tissue-patterning machinery, the field-mediated bioelectric model introduced previously (Manicka and Levin 2025). Each embryo is represented as a two-dimensional lattice of cells. Every cell carries a pair of voltage-gated ion channels, one depolarising and one polarising, whose joint action makes the cell bistable, so that its membrane potential settles to one of two stable values separated by an unstable equilibrium. Neighbouring cells are connected by gap junctions whose conductance depends on the difference in membrane potential across them, being highest between cells at similar potential and lowest across a steep difference, which allows the tissue to hold spatially distinct regions rather than relaxing to a uniform state. Superimposed on this lattice is an intra-embryonic electric field, aggregated from every cell’s charge, that feeds back on the conductance of the polarising channel: how strongly it can do so is set by each cell’s own biochemical network, driven by that cell’s membrane potential and, at the embryo level, by the field-shifted damping above, through an intermediate signal the network produces. A cell’s response to its field environment is therefore not a fixed parameter but a dynamic variable, set by the biochemical state the network has reached.

At the end of the simulated developmental window, the pattern of membrane potential this machinery has produced across the tissue is compared to the pattern characteristic of normal development, and the embryo is scored as rescued if the deviation between the two falls below a fixed threshold. Whether an embryo is rescued is therefore set jointly by where it sits in the collective field and by the resilience it happens to have been assigned.

### Model equations

Emission is constitutive rather than triggered by rescue itself: a scheme in which embryos emit only after sensing a rescue signal would require that signal to already exist before emission begins, a circularity that constitutive emission avoids.

Each embryo’s stress *Si* evolves independently of its bioelectric state as

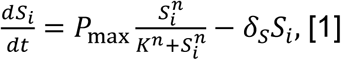

where *n* = 2 gives Hill-cooperative self-activation opposed by a fixed linear decay, so *Si* settles to one of two stable states, a low state near zero and a high, sustained one, separated by an unstable separatrix, the threshold value of *Si* below which it decays back to the low state and above which it commits to the high one: *S*^∗^ = 0.134 at the fitted parameters *P*max = 0.2/s, *K* = 0.5, *δS* = 0.1/s. Eq. 1 is stepped *k* = 500 times per bio-step, reflecting a separation of timescales between fast intracellular biochemistry and the slower tissue-level patterning it drives. Initial stress *Si*(0) is drawn independently per embryo per trial from a normal distribution with mean 0.3 and standard deviation 0.1, truncated below at *S*^∗^ so that every simulated embryo begins already past the point of commitment to the stress response, consistent with modelling a population already exposed to the teratogen at simulation onset.

The extracellular field *F* (eATP/adenosine; *μ*M) evolves as

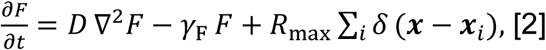

with absorbing boundary conditions *F* = 0 at the group perimeter. Here *D* = 11,500 *μ*m^2^/s is the eATP diffusivity, inferred from the injury-wave propagation speed (Tung, Sperry et al. 2024); *γ*F = 1.845 × 10^−4^/s i s the first-order decay rate (ectonucleotidase activity). Each embryo’s true emission rate is time-varying, *R*max*Si*(*t*), gated by its own stress (Eq. 1); *R*max is the per-embryo emission rate, calibrated so the boundary-free level *P*/*γ*F = 0.084 *μ*M matches the peak measured eATP concentration of 84 nM (Tung, Sperry et al. 2024); and the sum runs over all embryo positions on a square lattice with spacing ℎ = 3,000 *μ*m. The steady-state profile below is reached once every embryo’s stress has saturated, which the simulation confirms happens well before the run ends (SI Appendix); along the group midline it is

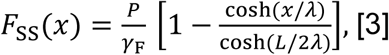

where *x* = 0 is the group center, *x* = ±*L*/2 are the absorbing boundaries, 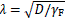 is the signal decay length, and *P* = *R*max/ℎ^2^ is the volumetric emission density.

Two consequences of Eq. 3 follow directly for how this steady state changes with group size. At the centre (*x* = 0),

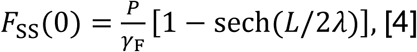

where *P*/*γ*F is the concentration the field would reach with no boundary at all, and sech(*L*/2*λ*) is the fraction of that concentration the boundary still withholds: near unity for

*L* ≪ *λ*, falling toward zero for *L* ≫ *λ*. At the boundary (*x* = *L*/2), the gradient is

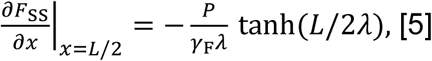

which converges to the fixed value *P*/(*γ*F*λ*), independent of *L*, once *L* ≫ *λ*. Growth therefore does not weaken the boundary’s local pull; it confines that pull to a rim of fixed width set by *λ*, while the fraction of the group that sits within this rim shrinks as the group grows. Checked against the simulated field at each group’s centre (SI Appendix 1), Eq. 4 reproduces the correct saturation shape and ceiling but over-predicts the absolute concentration by a roughly constant factor of ∼ 2 for *N* ≥ 25, chiefly because the production field’s distance-weighted coupling and boundary leakage remove more signal than the idealised uniform lattice.

The field feeds back on embryo *i*’s biochemical network by shifting its effective damping,

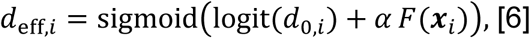

where logit(*p*) = ln(*p*/(1 − *p*)) is the log-odds transform, *d*_0_,*i* is embryo *i*’s intrinsic resilience, an independently assigned baseline capacity to reach the normal pattern without the field’s help, and *α* = 400 is the bioelectric coupling gain. This shift scales the biochemical network’s own connection weights multiplicatively (each weight is multiplied by *d*eff,*i*), biasing the network’s output and, through it, each cell’s field-sensitivity (below); at the tissue level this biases the whole embryo toward its normal developmental attractor, the coordinated membrane-potential pattern characteristic of normal development (as opposed to each individual cell’s own bistable choice between the two stable values described above), as *d*eff,*i* rises. Values of *d*0,*i* are drawn independently per embryo per trial from 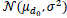 with *μd* = 0.01 and *σ* = 0.040, then clipped to the valid range [0.001,0.999]. Because the lower bound lies only 0.225*σ* below the nominal mean, clipping is not a negligible correction at these parameters: 41% of draws are pinned at the floor *d*_0_ = 0.001 and the realised mean is 0.022 rather than the nominal 0.010. The resulting distribution is therefore bimodal, with a point mass of highly susceptible embryos alongside a continuous tail of more resilient ones, and all reported results use this clipped distribution.

An approximate, illustrative field threshold and critical group size, neither used to score rescue in this paper (below), can be derived from an isolated embryo’s own field-driven bifurcation point (SI Appendix 6).

Within the bioelectric model, each cell’s membrane potential *V* evolves as

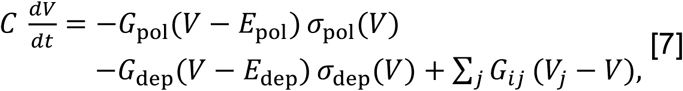

where *σ*pol and *σ*dep are voltage-gating sigmoids for the polarising and depolarising channels respectively, and the gap-junction conductance *Gij* between neighbouring cells *i*, *j* itself depends on their voltage difference, falling as it steepens. *G*dep is fixed; *G*pol is not, and evolves by its own dynamics (below). Full derivation of the channel and gap-junction dynamics, and of the fitted constants *G*dep, *E*pol, *E*dep, *C*, is in (Manicka and Levin 2025).

The intra-embryonic field mentioned above aggregates every cell’s charge *Qc* = *CVc* at each field point *p*,

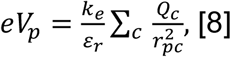

a Coulomb’s-law sum over cells *c* at distance *rpc* from *p* (shown here as a scalar magnitude; the full two-dimensional vector form, and the fitted constants *ke*/*εr*, are in (Manicka and Levin 2025)).

The biochemical network mentioned above is a set of interacting species ***η*** within each cell, driven by that cell’s own membrane potential and, at the embryo level, by the field-shifted damping *d*eff,*i* (Eq. 6):

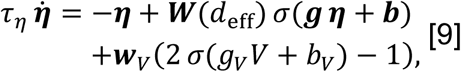

where ***W***(*d*eff) is the network’s own within– and between-cell interaction matrix, scaled by *d*eff,*i* above, and *τη*, ***g***, ***b***, ***w****V*, *gV*, *bV* are fitted constants (six interacting species per cell; full connectivity in (Manicka and Levin 2025)).

This network’s only output in the fitted configuration used here is a per-cell ligand concentration ℓ, which it drives and which also diffuses between neighbouring cells *j*,

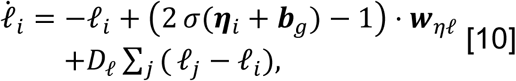

where ***w****η*ℓ is the network-to-ligand weight vector and *D*ℓ is a fitted diffusion strength; the network has no separate route back to *V* in this configuration; every effect it has on the tissue’s electrical dynamics runs through ℓ. This ligand concentration in turn sets each cell’s sensitivity to the local field,

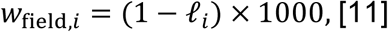

which finally closes the loop, gating how strongly the local field *eVi* (averaged over cell *i*’s neighbourhood) can shift *G*pol,

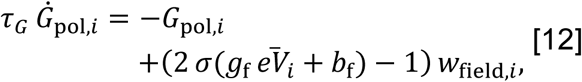

with *τG*, *g*f, *b*f fitted constants ((Manicka and Levin 2025)): at high ℓ*i*, *w*field,*i* falls toward zero and the cell is desensitised to the field regardless of the field’s own strength.

At the end of the run, embryo *i*’s Vmem pattern is compared to the reference pattern characteristic of normal development,

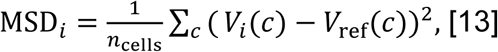

and embryo *i* is scored as rescued if MSD*i* < 2.33 × 10^−4^ V ^2^, the criterion used throughout this paper unless otherwise noted.

### Biological parameter calibration

The model is minimal in a specific sense: only four quantities are fit to the observed rescue curve, two describing the shared field (*γ*F, *α*) and two describing individual embryos (*σ*, the spread of intrinsic resilience, and the MSD rescue threshold). The remaining constants appearing in Eqs. 1–6 are either anchored to independent measurements, characterised in a separate experiment, or chosen on structural grounds, summarised below.

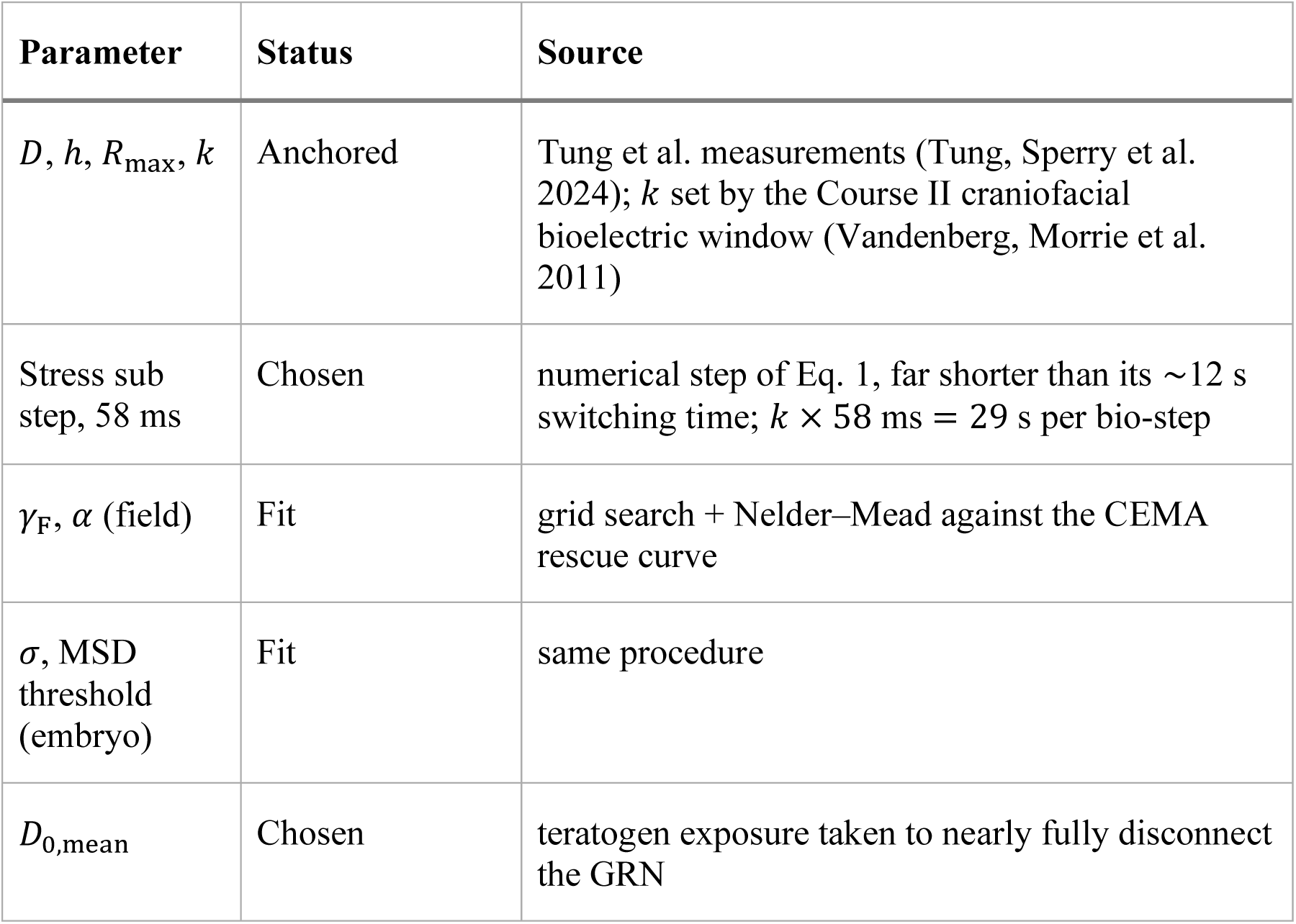
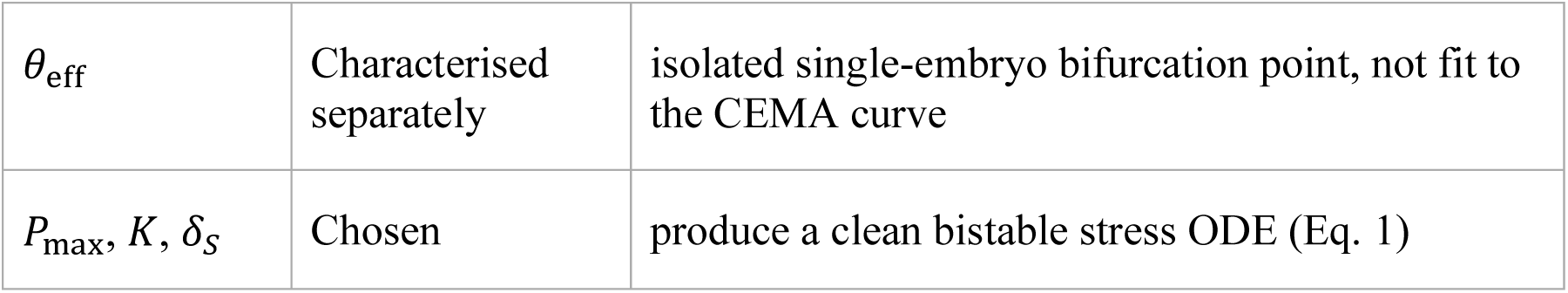

The fitted decay rate *γ*F (half-life ≈ 63 min) and coupling gain *α* fall within the broad range of timescales and sensitivities documented for extracellular purinergic signalling generally: release, ectonucleotidase-mediated degradation, and receptor engagement typically proceed on a timescale of minutes to tens of minutes (Burnstock 2007). *Xenopus* embryos express the relevant ectonucleotidase and purinergic receptor machinery at the developmental stages studied here (Blanchard, Boué-Grabot et al. 2019, Massé, Bhamra et al. 2021). We do not attempt to resolve which specific receptor subtype or downstream cascade step is operative; the fit constrains the combination *γ*F/*α*, not the underlying pharmacology (Discussion).

### Model simulation

Simulations resolved every embryo and coupled them through the shared field. A group of *N* embryos occupies the nodes of a rectangular lattice with spacing ℎ = 3,000 *μ*m, and each nominal group size was mapped to the nearest compact rectangle: the six experimental sizes *N* = 1,5,25,75,100 and 300 correspond to grids of 1 × 1, 2 × 3, 5 × 5, 9 × 9, 10 × 10 and 17 × 18 embryos (1, 6, 25, 81, 100 and 306 embryos), and the analyses of field structure and shuffling add 6 × 6, 7 × 7, 8 × 8 and 14 × 15 grids. Each embryo is an 11 × 11 lattice of cells (121 cells, six intracellular species each) running the dynamics of Eqs. 7–12.

The field *F* is represented at the embryo positions rather than on a finer mesh. Every pair of embryos within five decay lengths exchanges signal with weight *D* Δ*t*/*d*^2^, where *d* is their separation and Δ*t* the duration of one bio-step. Absorbing boundaries enter as sink terms of the same form for the lattice neighbours a boundary embryo lacks, which is equivalent to holding those neighbours at *F* = 0 (Eq. 2). A reflecting side, used in the boundary-control experiment, has no sink, which is equivalent to a mirror-image neighbour and enforces zero flux. Diffusion, decay and emission were integrated by explicit Euler steps, with at least ten sub-steps per bio-step and more whenever needed to keep the Courant number below one.

Each bio-step advances the stress variables (Eq. 1), integrated in *k* = 500 Euler sub-steps of 58 ms, then the field, and then every embryo’s network under its updated effective damping (Eq. 6). One bio-step therefore spans 29 s of biological time, and *k* was set so that a run of 1,000 bio-steps (about 8 h) covers the craniofacial bioelectric patterning window (Vandenberg, Morrie et al. 2011). Rescue was scored on the final Vmem pattern (Eq. 13) against a reference obtained by running an unperturbed embryo, with an effective damping of one and no field, through the same protocol. Further details of the morphogenetic patterning process are given in (Manicka and Levin 2025).

Trials differed in the independent per-embryo draws of *Si*(0) and *d*0,*i*. Group-size sweeps used 50 trials per size, the shuffle experiments 50 trials per condition, and the boundary-control experiments 10 per configuration.

### Shuffle experiments and noise-ceiling normalisation

Embryos were permuted to new grid positions at bio-step *t*shuffle ∈ {0,1,5,20,50,100} and the simulation continued to completion; 50 trials per condition. Two quantities were computed. The *positional* metric is the Pearson *r*^2^ between the trial-averaged spatial MSD map of the shuffled run and that of the unshuffled baseline, both indexed by grid position. The *identity* metric places each embryo’s final MSD at its own pre-shuffle position and correlates the result with the baseline within each trial before averaging across trials, so that it reports whether the same embryo fares similarly regardless of where it is moved.

Because a trial-averaged map is itself reproducible only up to a noise floor, and that floor rises with *N* as the field flattens, the positional metric is reported relative to a per-*N* ceiling. The 50 baseline trials are split into two halves of 25, each half averaged, and the two averages correlated; the Spearman–Brown correction ceiling = [2*r*half/(1 + *r*half)]^2^ then gives the *r*^2^ expected between two independent 50-trial averages of the same position-indexed map. The ceiling falls from 0.89 at *N* = 25 to 0.56 at *N* = 306. Reported ratios are observed *r*^2^ divided by this ceiling, so that unity denotes reconstitution as complete as the data permit. Two features of the construction should be noted. Ratios may exceed unity because the shuffled runs reuse the baseline field realisation and therefore share a source of variance that the disjoint-halves ceiling excludes, which makes the ceiling conservative. Ratios are computed from deterministic mean-versus-mean correlations, which run somewhat higher than bootstrap-resampled means, the discrepancy growing with *N*; unnormalised *r*^2^ values and normalised ratios are therefore not directly comparable and are not quoted together.

Statistical significance of the erosion at t=20 and t=100 relative to t=0, within each N, was assessed by a paired bootstrap. Because baseline and shuffled-t trials are matched one-to-one by trial index, each of 2000 bootstrap draws resampled the baseline trial indices once and reused that resample, together with a freshly split-half ceiling computed from it, for both conditions being compared; this shares the ceiling-estimation noise between the two ratios and isolates the difference attributable to the shuffle itself. The two-sided p-value is twice the smaller of the fraction of bootstrap differences at or below zero and the fraction at or above zero. Benjamini–Hochberg false-discovery-rate correction was applied jointly across all fourteen tests (N ∈ {25, 36, 49, 64, 100, 210, 306} × {t=20, t=100}); asterisks in Fig. 7 mark q<0.05 (∗), q<0.01 (∗∗), and q<0.001 (∗∗∗).

## Data Availability

Simulation code and data will be made available at GitHub repository upon publication.

## Acknowledgments

This research was supported by Emerald Gate Charitable Trust and we thank members of the Levin lab for helpful discussions.

## AI Use

S.M. used AI tools to partly assist with coding, analysis and manuscript writing. S.M. thoroughly reviewed and verified their output before incorporating them into the research pipeline. S.M. takes full responsibility for the accuracy and integrity of whatever AI-generated material that made its way into this paper.

## Supplementary Information

## Appendix 1: Derivation and validation of the closed-form centre-field prediction (Eqs. 3–4)

**Derivation.** Methods reduces the governing field equation (Eq. 2) to a 1D line-source idealisation along the group’s midline: the discrete embryo emitters are replaced by a continuous, uniform volumetric source *P* = *R*max/ℎ^2^ spread over −*L*/2 < *x* < *L*/2, with 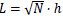 (the group modelled as a square of side length 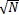 lattice spacings) and absorbing boundaries at the group edge. At steady state this is the boundary value problem

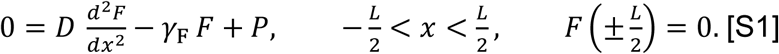

Writing 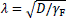 (Eq. 3’s decay length) and dividing through by *D* gives *F*^″^ − *F*/*λ*^2^ = −*P*/*D*. The homogeneous equation 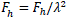 has general solution

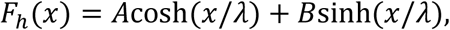

and a constant *Fp* = *P*/*γ*F solves the inhomogeneous equation directly, so the general solution is

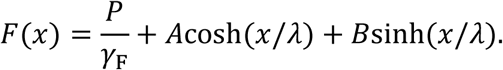

The source and the boundary conditions are both symmetric under *x* → −*x*, so *F* must be even; since sinh is odd and cosh is even, this forces *B* = 0. The remaining boundary condition, *F*(*L*/2) = 0, then fixes *A*:

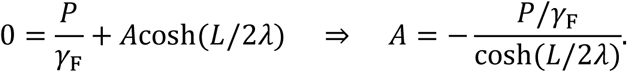

Substituting *A* back gives the general solution along the midline,

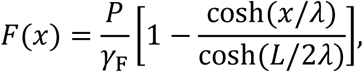

which is Eq. 3. Setting *x* = 0, where cosh(0) = 1, gives the centre-field special case

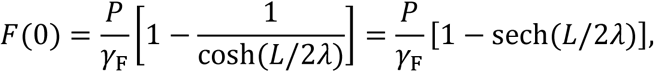

which is Eq. 4.

**Validation.** We checked Eq. 4 against the simulated field’s centre value, averaged over the same 50-trial runs behind Fig. 2a, at every canonical group size, and against an intermediate 2D uniform-absorbing-lattice proxy that shares Eq. 4’s idealised, distance-independent coupling but is solved on the actual 2D grid rather than reduced to 1D.

**Figure S1.**
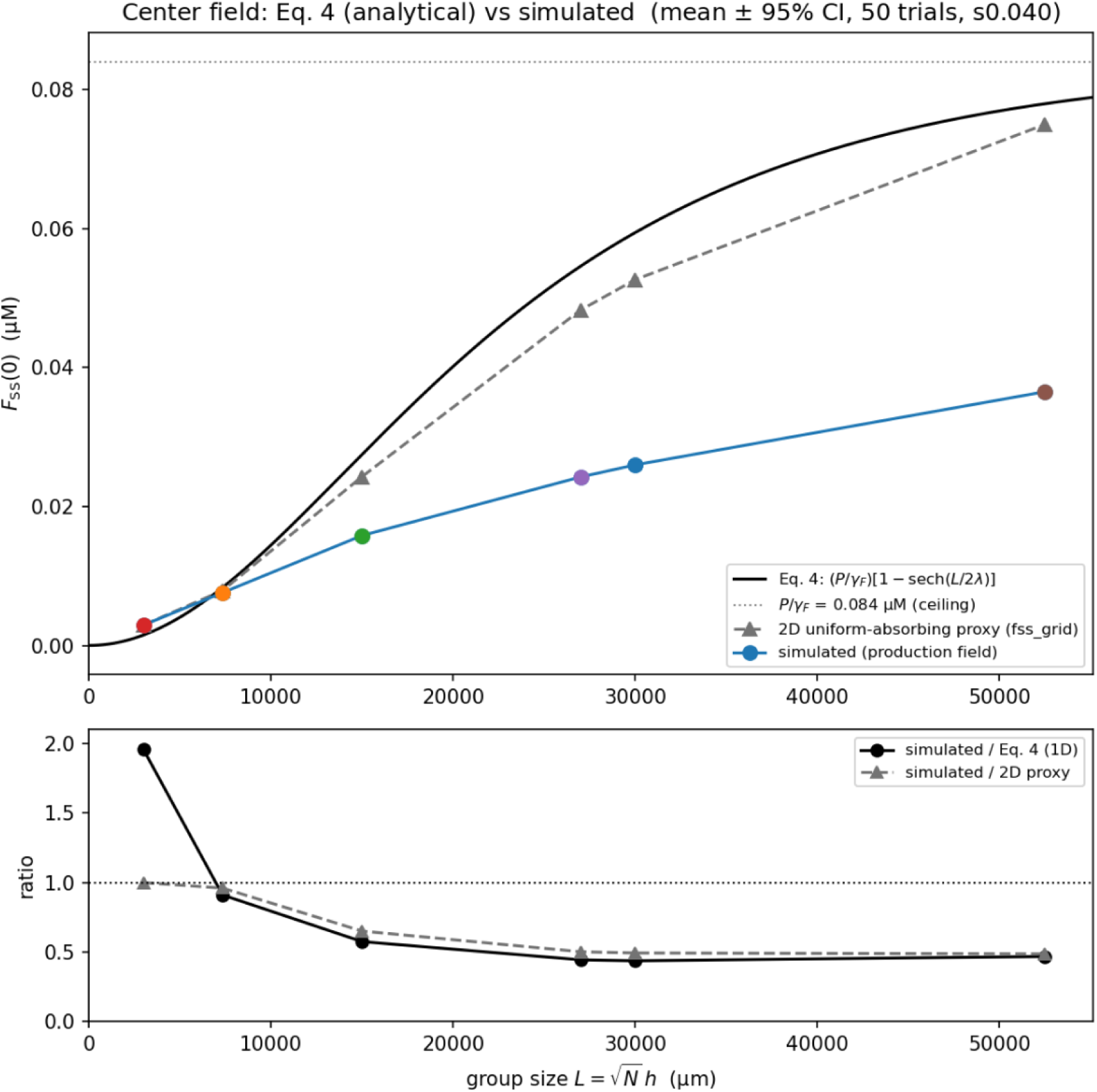
Eq. 4 reproduces the correct saturation shape and ceiling but over-predicts the simulated centre field by a roughly constant factor of ∼2 for *N* ≥ 25. Top: analytical prediction (Eq. 4, black), the 2D uniform-absorbing proxy (grey triangles), and the simulated production field (coloured circles; mean of 50 trials, 95% CI smaller than the marker since the final-step centre field is deterministic across trials) against group size *L* = √*N* ℎ. Bottom: the ratio of simulated to each reference curve.

| *N* | grid | *F*<sub>sim</sub> (*μ*M) | *F*<sub>Eq.4</sub> (*μ*M) | sim/Eq.4 | sim/2D |
|---|---|---|---|---|---|
| 1 | 1×1 | 0.00293 | 0.00149 | 1.959 | 1.000 |
| 6 | 2×3 | 0.00760 | 0.00834 | 0.911 | 0.961 |
| 25 | 5×5 | 0.01582 | 0.02748 | 0.576 | 0.651 |
| 81 | 9×9 | 0.02425 | 0.05458 | 0.444 | 0.502 |
| 100 | 10×10 | 0.02599 | 0.05942 | 0.437 | 0.494 |
| 306 | 17×18 | 0.03651 | 0.07796 | 0.468 | 0.487 |

Eq. 4 correctly locates the boundary-free ceiling, *P*/*γ*F = 0.084 *μ*M, matching the 84 nM calibration target (Methods), and reproduces the qualitative saturation shape: both curves rise from near zero and flatten toward the same ceiling as *N* grows. It over-predicts the simulated centre field by a roughly constant factor of ∼2 for *N* ≥ 25 (sim/Eq.4 ratio 0.44–0.58, plateauing at 0.44–0.47 for the largest groups). The gap decomposes into two parts: about 12% is attributable to the 1D→2D idealisation alone (the sim/2D column is closer to unity than sim/Eq.4 at every *N*), and the remaining, larger and growing-with-*N* share (35–51%) comes from the production field’s distance-weighted adjacency and explicit boundary leakage, both absent from the uniform-lattice proxies. *N* = 1 is a degenerate case (no group midline, *L* = ℎ), where the simulated field matches the 2D proxy exactly by construction.

## Appendix 2: Single-trial rescue maps at *N* = 25 and *N* = 100

Result 2 states that a single trial’s outcome is spatially disorganised near threshold but already centre-rescued well above it, before averaging over trials is introduced. Figure S2 shows the per-embryo MSD (Eq. 13) underlying that claim, for the same representative trials used in Fig. 3 (*N* = 25, 8/25 rescued; *N* = 100, 75/100 rescued), with the rescue threshold (MSD < 2.33 × 10^−4^ V^2^) overlaid as a dashed contour.

**Figure S2.**
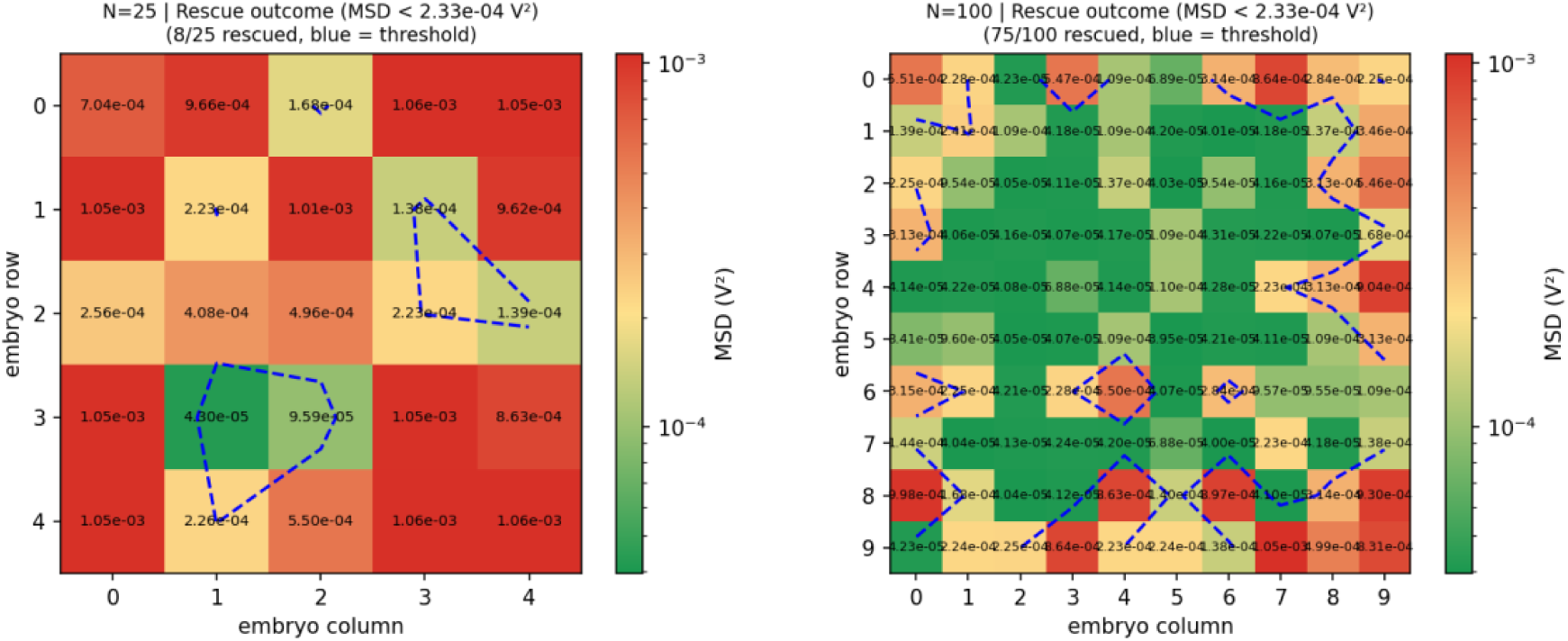
A single trial’s rescue outcome is spatially disorganised near threshold but already centre-rescued well above it. Per-embryo MSD (colour, log scale) for one representative trial at *N* = 25 (left) and *N* = 100 (right); the dashed contour marks the rescue threshold. At *N* = 25 rescued embryos (inside the contour) form no coherent region relative to the group centre. At *N* = 100 the rescued region is centred on the group and excludes the corners, even in this single, unaveraged trial.

At *N* = 25, only 8 of 25 embryos clear the threshold, and the rescued region is fragmented into two small, disconnected patches (row 3, columns 1–2; and row 1–2, columns 3–4) rather than one contiguous area: the group’s own geometric centre (row 2, column 2; MSD = 4.96 × 10^−4^) is itself above threshold and therefore not rescued. At *N* = 100, where 75 of 100 embryos clear the threshold, the rescued cells instead form one large, contiguous, interior-dominant region, with unrescued cells concentrated toward the perimeter, even though *d*_0_ is drawn independently per embryo and no trial-averaging has been applied. This is consistent with the centre-to-corner field gradient (61% at *N* = 25, 79% at *N* = 100; Result 2) being large enough, relative to *d*_0_’s spread, to fix the sign of most embryos’ outcomes by position alone at *N* = 100 but not yet at *N* = 25, where the field’s centre-to-edge contrast is too shallow for *F* to consistently outweigh *d*_0_ in single-trial outcomes. Result 2’s trial-averaged maps (Fig. 5) recover a centre-to-edge gradient at every group size, including *N* = 25, by cancelling this *d*_0_ noise rather than by any change in the field itself.

## Appendix 3: Isolated-embryo causal test of field dominance over intrinsic resilience

Result 2 reports a direct causal test that the field, rather than an embryo’s intrinsic resilience *d*_0_, determines rescue outcome, isolating a single embryo ( *N* = 1, no neighbours, no field diffusion) and externally fixing *d*_0_ and *F* independently, removing the correlation between them that exists in a real group. *d*_0_ was swept over seven values at two spacings (log– and linear-spaced, spanning its full sampled range, to check the result does not depend on this choice); *F* was fixed at the eight mean steady-state field values observed at the eight canonical group sizes in the main rescue sweep (Fig. 2). Each of the resulting 7 × 8 combinations is a single deterministic embryo, so the per-cell MSD (Eq. 13) needs no trial-averaging. Figure S3 shows the resulting MSD surface at both *d*_0_ spacings.

**Figure S3.**
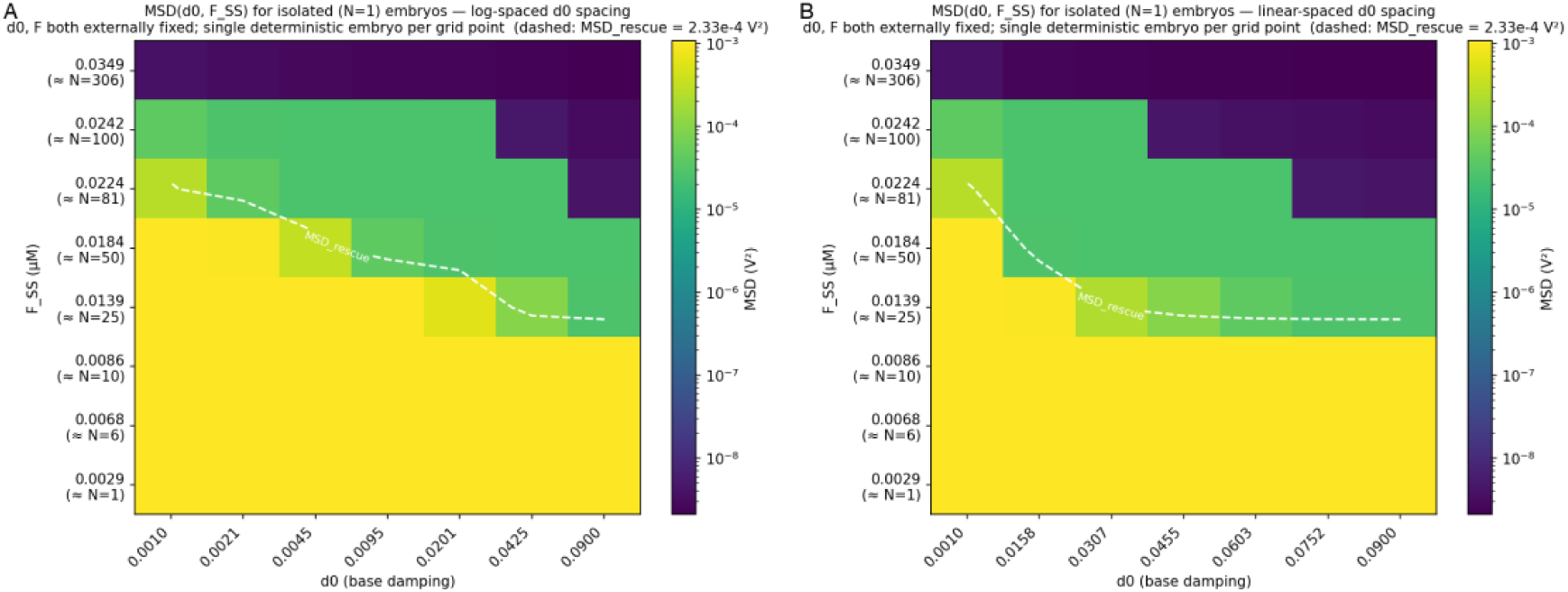
Fixing *F* alone determines rescue outcome; fixing *d*_0_ alone does not. Per-embryo MSD (colour, log scale) for an isolated embryo with *d*_0_ and *F* both externally fixed, at every combination of seven *d*_0_ values (log-spaced, **A**; linear-spaced, **B**) and the eight canonical field values from Fig. 2. The dashed contour marks the rescue threshold (*MSD* < 2.33 × 10^−4^ V^2^). Colour bands run horizontally (set by *F*) rather than vertically (set by *d*_0_) at every group size.

A two-way ANOVA decomposition of the variance in log(MSD) across this grid attributes 81.6% (log-spaced *d*_0_) or 79.4% (linear-spaced *d*_0_) to *F* and 6.4% to *d*_0_ in both cases (the remainder, 12.0–14.2%, is interaction/nonlinearity), matching the 79–82%/6.4% split reported in Result 2. At the two *d*_0_ extremes tested (0.001 and 0.09), fixing *F* at its lowest value gives log(MSD) = −6.85 for both, identically, and neither rescues; fixing *F* at its highest value gives −19.3 and −20.0 respectively, and both rescue: *F* alone fixes the outcome at either extreme, irrespective of *d*_0_. The converse does not hold: at every one of the seven *d*_0_ values tested, sweeping *F* across its full range still changes log(MSD) by 12.5–13.2, comparable to the full range seen when *d*_0_ is unconstrained, so no value of *d*_0_ pins the outcome irrespective of *F*.

## Appendix 4: A saddle-node bifurcation in the interior cells’ polarising-channel conductance selects fate

Fig. 4 attributes the divergence between rescued and failing patterns to a saddle-node bifurcation in *G*pol, the interior cells’ polarising-channel conductance. Dropping the gap-junction term in Eq. 7 (an isolated cell, no neighbours) and holding every other channel constant at its fitted value, the fixed points of *C dV*/*dt* = 0 depend on *G*pol alone; sweeping *G*pol and classifying each root by the sign of *∂*(*dV*/*dt*)/ *∂V* locates where a stable and an unstable fixed point collide and annihilate. Figure S4 shows this bifurcation diagram together with the actual interior *G*pol(*t*) trajectories of representative rescued and failing embryos at three group sizes.

**Figure S4.**
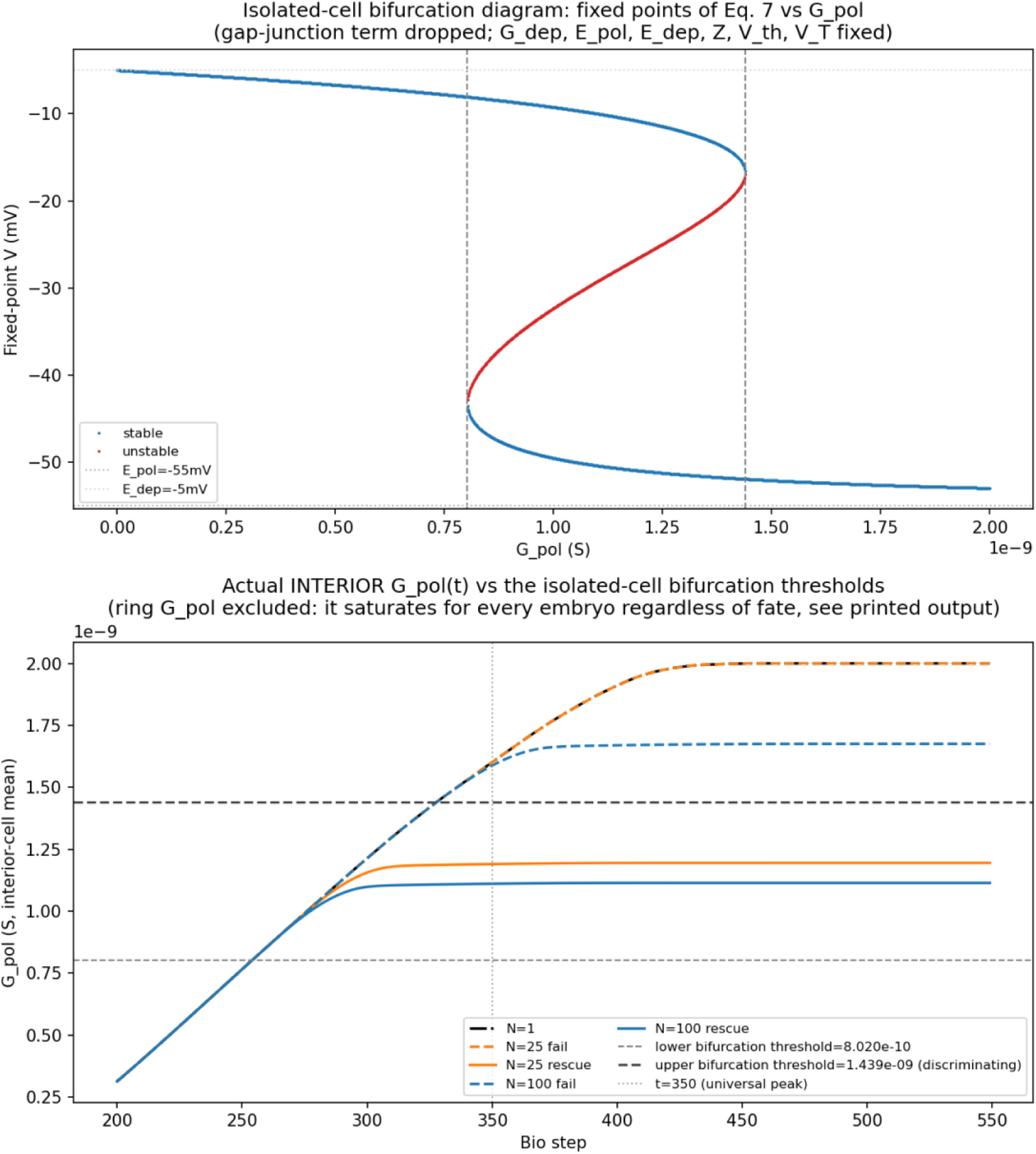
Rescued embryos stay inside the bistable window; failing embryos are driven out of it. **Top**: fixed points of the isolated-cell membrane equation (Eq. 7, gap-junction term dropped) as a function of *Gpol*, holding *Gdep* and ther channel constants fixed. Two stable branches (blue) are joined by an unstable branch (red); the two saddle-node thresholds where a stable and unstable branch meet are marked by dashed vertical lines. **Bottom:** interior-cell mean *Gpol*(*t*) (Eq. 12) for one rescued and one failing representative embryo at *N* = 1,25,100, against the same two thresholds.

The saddle-node thresholds are *G*pol = 8.02 × 10^−10^ S (lower) and 1.439 × 10^−9^ S (upper), bracketing a bistable window that contains every embryo’s shared initial value (*G*pol(0) = 1.0 × 10^−9^ S). Every embryo’s interior *G*pol rises together at first, tracking the field-gated ligand desensitisation common to the whole group (Eq. 12): this shared rise, common to every eventual outcome, is what produces the shared early MSD dip in Fig. 4. The two fates separate only once *G*pol approaches the upper threshold. Rescued embryos’ *G*pol plateaus below it and remains inside the bistable window, so the cell retains its normal hyperpolarised fixed point; failing embryos’ *G*pol instead continues past the upper threshold into the range where that hyperpolarised fixed point has already been annihilated, leaving only the depolarised branch. This is a genuine bifurcation in the underlying dynamics, not merely a descriptive threshold on *G*pol(*t*): past the upper threshold, the rescued state is not a nearby attractor the cell fails to reach quickly enough, but one that no longer exists.

## Appendix 5: The shuffle’s identity component

Fig. 7 reports the *positional* component of the shuffle perturbation: whether the trial-averaged rescue map is reconstituted at each grid position after relocating embryos at time *t*. A complementary *identity* component asks the opposite question of the same shuffled runs: remapping each shuffled embryo’s outcome back to its own original position (using the recorded permutation) and correlating that remapped outcome, embryo by embryo, against its own outcome in the matched unshuffled baseline. A Pearson *r* near 1 means an embryo’s fate stays tied to that embryo regardless of where the shuffle sent it (identity-tracking); *r* near 0 would mean fate is uncorrelated with identity once position is factored out. Figure S5 reports this identity correlation across the same shuffle times and an overlapping range of group sizes (*N* = 25–210).

**Figure S5.**
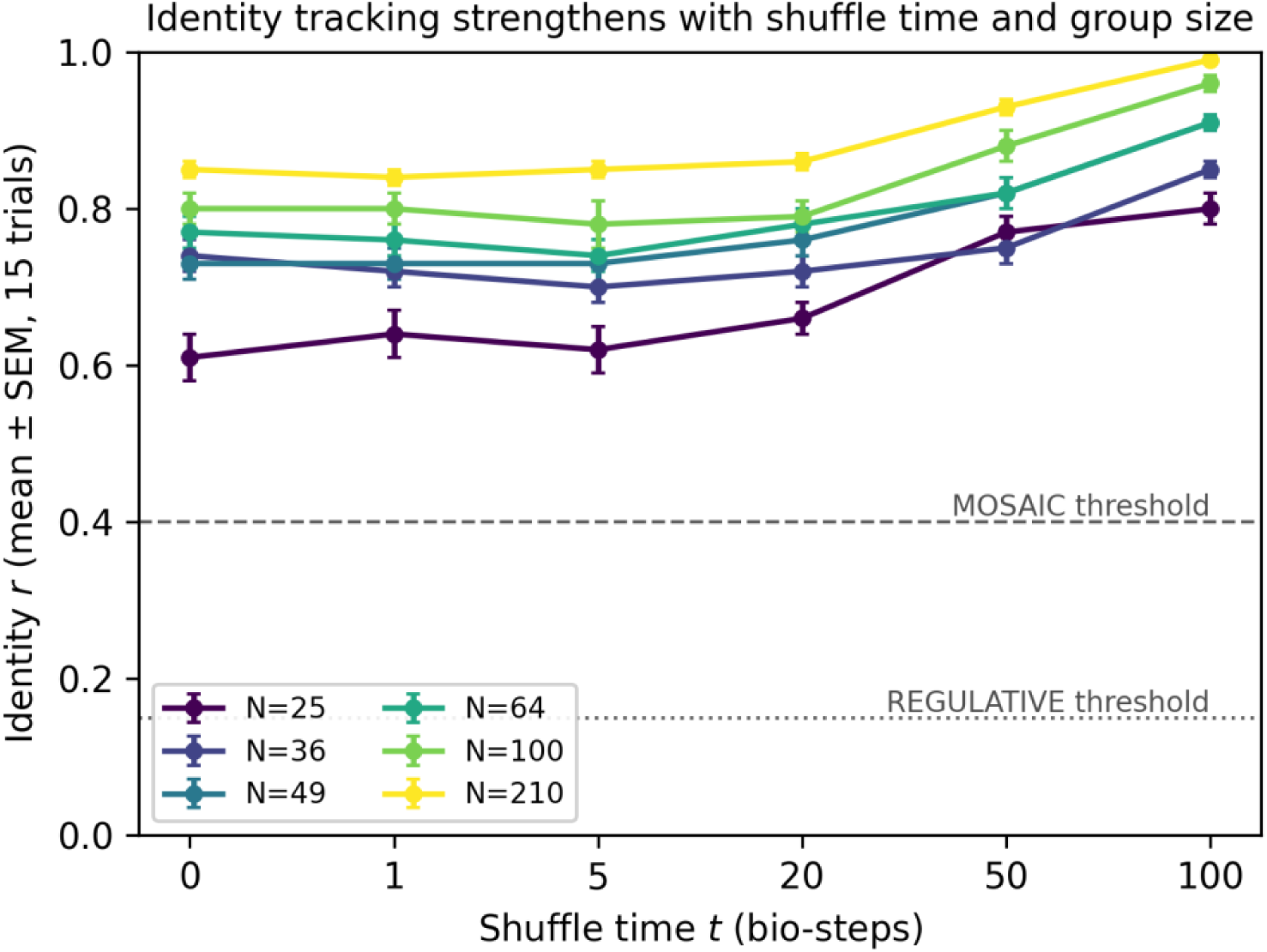
An embryo’s fate tracks its own identity, not the position the shuffle sent it to, and does so more strongly at later shuffle times and in larger groups. Per-trial Pearson *r* (mean ± SEM, 15 trials) between each shuffled embryo’s outcome, remapped to its original position, and that same embryo’s unshuffled-baseline outcome, plotted against shuffle time for each group size tested (*N* = 25–210). Dashed line: MOSAIC threshold (*r* ≥ 0.4, fate follows the embryo); dotted line: REGULATIVE threshold (*r* ≤ 0.15, fate follows the position). Every point lies in the MOSAIC range.

Identity *r* is MOSAIC-range (0.61–0.99) at every group size and shuffle time tested, in contrast to the positional ratio of Fig. 7, which collapses toward 0 by *t* = 100 at large *N*. Rather than declining with shuffle time, identity *r* rises with it at every *N*: at *N* = 25 it goes from 0.61 ± 0.03 at *t* = 0 to 0.80 ± 0.02 at *t* = 100; at *N* = 210, from 0.85 ± 0.01 to 0.99 ± 0.00. It also rises with group size at fixed *t* (e.g. at *t* = 100, from 0.80 at *N* = 25 to 0.99 at *N* = 210), the same direction as the ceiling-relative positional ratio in Fig. 7. Together with the positional result, this shows the two components of the shuffle diverge rather than move together: once the positional commitment window has closed, relocating an embryo no longer redirects its fate, but that fate has not become noise, it has simply reattached to the embryo itself, and does so more tightly, not less, as *N* grows.

## Appendix 6: The empirical bifurcation threshold *θ*eff and the approximate field threshold it implies

Methods points here for an approximate, illustrative rescue field threshold *F*^∗^ and critical group size *Lc*, derived below from *θ*eff, an isolated embryo’s own field-driven bifurcation point: the value of the effective damping *d*eff (Eq. 6) at which its tissue-level pattern commits to the normal attractor. We measured *θ*eff directly by forcing *d*0 to fixed values in an isolated embryo (*N* = 1, *d*_0_ and initial stress both deterministic, no other source of trial-to-trial variation) and running the full model at its canonical parameters (*α* = 400, *γ*F = 1.845 × 10^−4^/s, 1000 bio-steps), scoring rescue by the same MSD criterion used throughout (MSD < 2.33 × 10^−4^ V^2^).

Even an isolated embryo is not field-free: with no neighbours to receive a field from, it still constitutively emits into its own local medium (Eq. 2), and that self-field settles to a small steady state set purely by emission balanced against decay and boundary leakage. Because emission depends only on stress, which saturates to the same ceiling regardless of *d*_0_, this self-field is essentially constant across the sweep 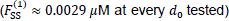, so it shifts every embryo’s effective damping by the same fixed amount in logit-space: 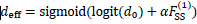 reproduces every measured *d*eff value from *d*0 alone.

## References

1. Allee, W. (1931). “Animal aggregations, a study in general sociology.,(The University of Chicago Press: Chicago, IL, USA).”

2. Bao, L., et al. (2004). “Pannexin membrane channels are mechanosensitive conduits for ATP.” FEBS Letters 572(1-3): 65–68.

3. Black, A. J., et al. (2020). “Ecological scaffolding and the evolution of individuality.” Nature Ecology & Evolution 4(3): 426–436.

4. Blanchard, C., et al. (2019). “Comparative Embryonic Spatio-Temporal Expression Profile Map of the Xenopus P2X Receptor Family.” Frontiers in Cellular Neuroscience 13.

5. Boedicker, J. Q., et al. (2009). “Microfluidic Confinement of Single Cells of Bacteria in Small Volumes Initiates High-Density Behavior of Quorum Sensing and Growth and Reveals Its Variability.” Angewandte Chemie International Edition 48(32): 5908–5911.

6. Burnstock, G. (2007). “Purine and pyrimidine receptors.” Cellular and Molecular Life Sciences 64(12): 1471–1483.

7. Burnstock, G. and H. Ulrich (2011). “Purinergic signaling in embryonic and stem cell development.” Cellular and Molecular Life Sciences 68(8): 1369–1394.

8. Clarke, E. (2013). “The multiple realizability of biological individuals.” The Journal of Philosophy 110(8).

9. Crick, F. (1970). “Diffusion in Embryogenesis.” Nature 225(5231): 420–422.

10. De Robertis, E. M. (2009). “Spemann’s organizer and the self-regulation of embryonic fields.” Mechanisms of Development 126(11-12): 925–941.

11. French, V., et al. (1976). “Pattern Regulation in Epimorphic Fields.” Science 193(4257): 969–981.

12. Gierer, A. and H. Meinhardt (1972). “A theory of biological pattern formation.” Kybernetik 12(1): 30–39.

13. Goodwin, B. C. and L. E. H. Trainor (1980). “A field description of the cleavage process in embryogenesis.” Journal of Theoretical Biology 85(4): 757–770.

14. Gurdon, J. B. (1988). “A community effect in animal development.” Nature 336(6201): 772–774.

15. Hense, B. A., et al. (2007). “Does efficiency sensing unify diffusion and quorum sensing?” Nature Reviews Microbiology 5(3): 230–239.

16. Hodgkin, A. L. and A. F. Huxley (1952). “A quantitative description of membrane current and its application to conduction and excitation in nerve.” The Journal of Physiology 117(4): 500–544.

17. Kierstead, H. and L. B. Slobodkin (1953). “The size of water masses containing plankton blooms.” Journal of Marine Research 12(1).

18. Kim, M. K., et al. (2016). “Local and global consequences of flow on bacterial quorum sensing.” Nature Microbiology 1(1): 15005.

19. Krakauer, D., et al. (2020). “The information theory of individuality.” Theory in Biosciences 139(2): 209–223.

20. Levin, M. (2011). “The Wisdom of the Body: Future Techniques and Approaches to Morphogenetic Fields in Regenerative Medicine, Developmental Biology and Cancer.” Regenerative Medicine 6(6): 667–673.

21. Levin, M. (2019). “The Computational Boundary of a “Self”: Developmental Bioelectricity Drives Multicellularity and Scale-Free Cognition.” Frontiers in psychology 10.

22. Levin, M. (2022). “Technological Approach to Mind Everywhere: An Experimentally-Grounded Framework for Understanding Diverse Bodies and Minds.” Frontiers in Systems Neuroscience 16: 768201.

23. Lisman, J. E. (1985). “A mechanism for memory storage insensitive to molecular turnover: a bistable autophosphorylating kinase.” Proceedings of the National Academy of Sciences 82(9): 3055–3057.

24. Liu, Y., et al. (2020). “Thioridazine Induces Cardiotoxicity via Reactive Oxygen Species-Mediated hERG Channel Deficiency and L-Type Calcium Channel Activation.” Oxidative Medicine and Cellular Longevity 2020: 1–17.

25. Manicka, S. and M. Levin (2025). “Field-mediated bioelectric basis of morphogenetic prepatterning.” Cell Reports Physical Science 6(10).

26. Massé, K., et al. (2021). “The enpp4 ectonucleotidase regulates kidney patterning signalling networks in Xenopus embryos.” Communications Biology 4(1).

27. McMillen, P. and M. Levin (2024). “Collective intelligence: A unifying concept for integrating biology across scales and substrates.” Communications Biology 7(1).

28. Redfield, R. (2002). “Is quorum sensing a side effect of diffusion sensing?” Trends in Microbiology 10(8): 365–370.

29. Sergeeva, E. G., et al. (2026). Living multicellular systems induce decodable spatial patterns in bacterial collectives. bioRxiv, openRxiv.

30. Skellam, J. G. (1951). “Random Dispersal in Theoretical Populations.” Biometrika 38(1-2): 196–218.

31. Stephens, P. A., et al. (1999). “What is the Allee effect?” Oikos: 185–190.

32. Tung, A., et al. (2024). “Embryos assist morphogenesis of others through calcium and ATP signaling mechanisms in collective teratogen resistance.” Nat Commun 15(1): 535.

33. Turing, A. M. (1952). “The chemical basis of morphogenesis.” Philosophical Transactions of the Royal Society of London. Series B, Biological Sciences 237(641): 37–72.

34. Vandenberg, L. N., et al. (2011). “V-ATPase-dependent ectodermal voltage and ph regionalization are required for craniofacial morphogenesis.” Developmental Dynamics 240(8): 1889–1904.

35. Varela, F. G., et al. (1974). “Autopoiesis: The organization of living systems, its characterization and a model.” Biosystems 5(4): 187–196.

36. Waddington, C. H. (1942). “Canalization of Development and the Inheritance of Acquired Characters.” Nature 150(3811): 563–565.

37. Waters, C. M. and B. L. Bassler (2005). “QUORUM SENSING: Cell-to-Cell Communication in Bacteria.” Annual review of cell and developmental biology 21(1): 319–346.

38. Wen, A. X., et al. (2026). “Antibiotics stimulate protein transfer to persister cells.” Science 392(6805).

39. Wolpert, L. (1969). “Positional information and the spatial pattern of cellular differentiation.” Journal of Theoretical Biology 25(1): 1–47.

